# Acute hypercapnic stress modulates innate and learned defensive behavior

**DOI:** 10.64898/2026.06.18.733090

**Authors:** Nesrine Merabet, Clara Guiraud, Pascal Züfle, Margaux Vieu, Clara Zirah, Anh-Kiet Truong, Carlotta Martelli, Emmanuel Perisse

## Abstract

Survival depends on avoiding threats, a process shaped by experience and internal states. Notably, acute stress can induce analgesia, yet the neural mechanisms by which stress alters aversive value coding remain unclear. Using *Drosophila*, we show that prior noxious experience induces intensity-dependent analgesia and triggers the release of CO_2_, a known stress signal. Surprisingly, this effect is mediated by tracheal dendrite (td) neurons in the respiratory system rather than classic olfactory pathways. We then demonstrated that activation of td neurons induces analgesia, whereas their inhibition suppresses it and restores normal nocifensive and learned behavior. Finally, high CO_2_ exposure decreases dopaminergic neuron responses to electric shocks, thereby impairing aversive memory formation. Together, we propose that under hypercapnic (high CO_2_) stress, td neurons modulate nociceptive computation and aversive value coding in the brain to facilitate appropriate innate and learned behavioral responses.

## INTRODUCTION

Avoiding threat is a primary behavior necessary to survive. As the environment dynamically changes, animals must integrate external sensory cues with internal state and past experience to learn and adjust their behavioral response towards noxious stimuli (*1–5*). Notably, acute stress can temporarily and adaptively decrease response to noxious stimuli, a phenomenon known as stress-induced analgesia (*6–9*). As a consequence, acute stress can alter the encoding of aversive signals, thereby altering future innate and learned behavior (*10–14*). Acute stress can arise from interoceptive information (*15*) and/or come from the environmental cues (*16*), such as stress pheromones (*17*, *18*). However, the neural structures and mechanisms responsible for the perception and integration of acute stress to modulate aversive signals and drive appropriate behavior remain elusive.

As most organisms, fruit flies innately avoid nociceptive stimuli such as shock, heat, acid or bitter (*19–22*). When paired with another sensory stimulus during aversive associative learning, such as pairing an odor with electric shocks, flies can assign a negative value to the odor thereby influencing future learned behavior. As in vertebrates (for review see (*23*, *24*)), punishment dopaminergic neurons (DANs), targeting the mushroom body (MB), receive and respond to nociceptive signal intensities (e.g. electric shock or heat) (*5*, *25–28*). These signals induce intensity-dependent plasticity in the MB after olfactory learning, promoting appropriate learned odor-avoidance (*5*). Additionally, flies can assign a relative “better than” value to the least aversive of two sequential olfactory experiences, via interactions between the punishment and reward systems in the MB network (*5*). In such a senario, a strong and noxious olfactory experience (i.e. odor A paired with 60V electric shock) is followed by a weaker (and better) one (i.e. odor B paired with 30V electric shock), recruiting specific sets of reward DANs (*5*). A strong and noxious olfactory experience (i.e. odor A paired with 60V electric shock) can therefore trigger acute stress (*15*, *29*) and induce analgesia (*8*). Thus, the encoding of absolute and relative aversive values is likely modulated by a prior stressful experience.

In response to stress/danger, most animals (and plants), emit alarm signals (for review see (*18*)). These signals can be of different modalities such as visual, auditory, tactile and/or olfactory to alert potential threats to others (conspecific or predator), but can also benefit the individual itself by adjusting its behavior. For instance, volatile alarm pheromones are perceived by olfactory or gustatory pathways to mostly induce avoidance behavior (*30*). Electric shock or mechanical stimulation quickly induce the production of social *Drosophila* stress odorant (dSO), of which CO_2_ is a major component that flies usually avoid (*31*). However, whether stressful CO_2_ can induce analgesia and alter the encoding of aversive signals to modulate innate and learned response to nociceptive stimuli remains unknown.

Here, we combined behavioral analysis, gas chromatography/mass spectrometry, genetics, neuronal manipulation, anatomical characterization, and *in vivo* two-photon calcium imaging to study the effect of prior noxious/stressful experience on nocifensive behavior and aversive value coding. We first found that, as in mammals, exposure to noxious stimulation induces a transient and intensity-dependent analgesic state that reduces fly’s avoidance to subsequent (shock/heat) aversive stimuli. Moreover, shock stimulation elevates CO_2_ levels in the fly’s environment in an intensity-dependent manner. Pre-exposure to CO_2_ mimics the effect of noxious stimulation by inducing analgesia to mild but not high shocks, and this does not occur through the CO_2_ olfactory pathway. Instead, we found that tracheal dendrite (td) neurons in the respiratory system relay hypercapnia (high CO_2_ in the organism) to induce analgesia. Gr28b.c expressing td neuron’s output target the central nervous system where it likely interacts with the nociceptive pathway. In line with this hypothesis, we show that high CO_2_ suppresses low but not high punishment DANs responses to low but not high shock stimulation, matching behavioral data. Together, our findings support a model where noxious stimuli trigger hypercapnic stress detected in the respiratory tracheal system to induce analgesia and modulate nociceptive signal integration, thereby altering aversive value coding and learned/innate defensive behavior.

## RESULTS

### Prior shock experience decreases aversive learning and avoidance of aversive stimuli

Previous work including ours showed that flies can learn to associate a sequence of different odors paired with different intensities of electric shocks (*5*, *32*). As stress can interfere with memory processing (*13*, *14*), a prior aversive experience can therefore interfere with a following one. We therefore first investigated the effect of prior aversive experience on learning performances in an olfactory associative assay. In a tube covered with copper wires, groups of flies were exposed to an odor A alone followed by an odor C paired with a 30V electric shock (Fig. 1A). In between these two experiences flies were exposed to either nothing or different types of aversive experiences: odor B or electric shocks alone, or odor B paired with electric shocks (Fig. 1A). The presentation of the odor B alone did not affect immediate memory performances during the choice between the odors A vs C in a T-maze (Fig. 1A). Moreover, pre-exposing flies to an aversive odor alone does not change future shock or odor avoidance (fig. S1, A and B) (*33*). However, presenting electric shocks alone or paired with the odor B significantly reduced memory performances proportionally to shock intensity (Fig. 1B). These results demonstrate that nociceptive, and not just unpleasant, prior aversive experience alters following shock and/or odor encoding during learning to reduce immediate memory performances.

**Fig. 1.**
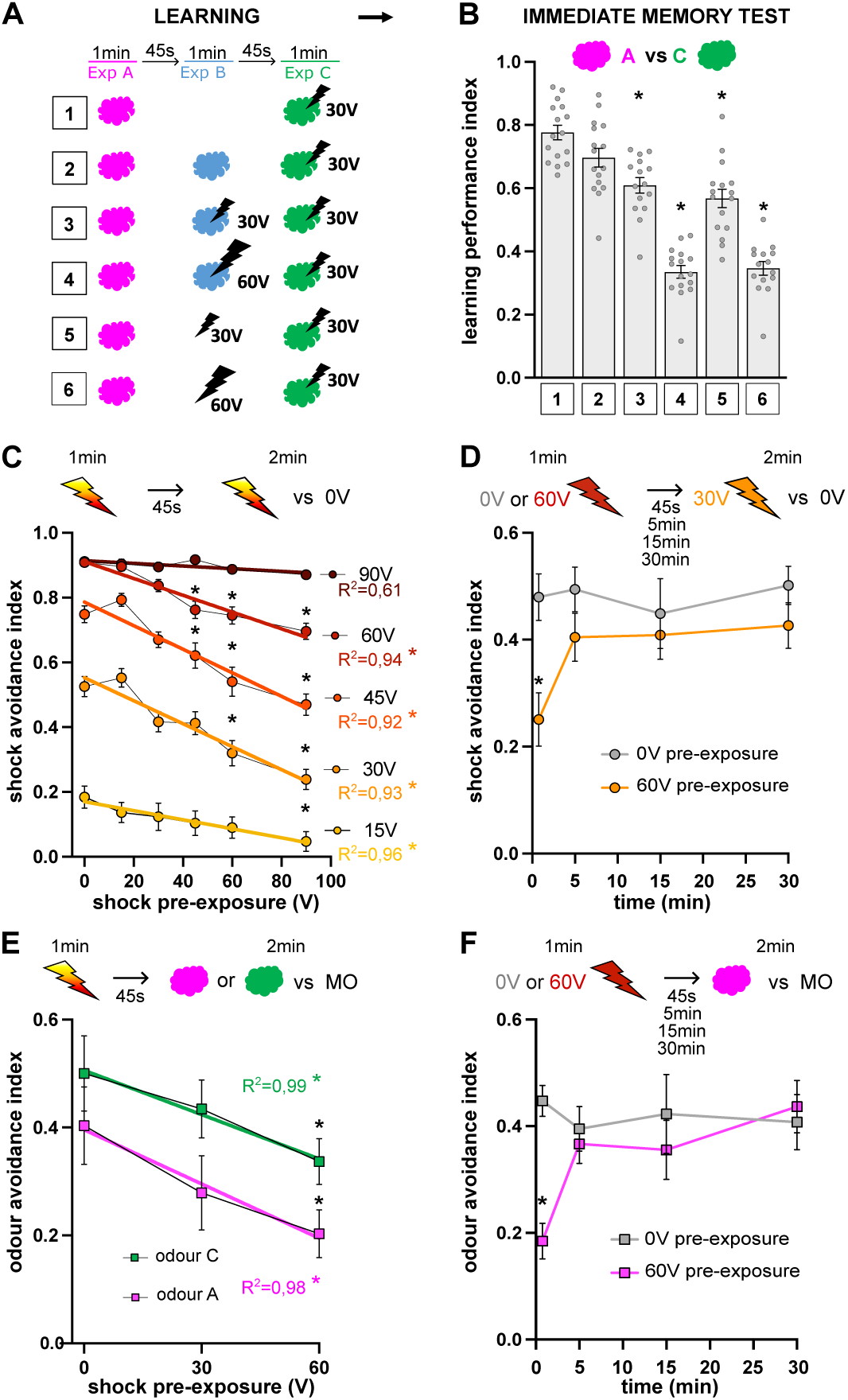
Prior shock experience decreases aversive learning and avoidance of aversive stimuli. (**A**) Immediate aversive olfactory learning protocol. (**B**) Immediate memory test performance index of odor A vs C significantly decreases with the strength of shock pre-exposure. Experimental groups (2-3-4-5-6) are statistically compared with the control group (1), n=15-16. (**C**) Shock avoidance index of 15V, 30V, 45V or 60V linearly decreases after shock pre-exposure but not for 90V shock avoidance, n=28-30. (**D**) Decrease of shock avoidance index (30V) after 60V shock pre-exposure comes back to normal after less than 5 min, n=12. (**E**) Odor avoidance index (OCT or MCH vs mineral oil (MO)) linearly decreases after (30V or 60V) shock pre-exposure, n=17-19. (**F**) Decrease of odor avoidance index after 60V shock pre-exposure comes back to normal after less than 5 min, n=11-12. Top of each graph represents the protocol. Data presented as mean ± standard error of mean (SEM).

We then asked whether a prior shock experience can alter future shock avoidance. We pre-exposed flies to a range of electric shock voltage intensities and then measured in a T-maze their avoidance of subsequent higher, similar, or lower voltage shock intensities. We found a strong and linear correlation between shock pre-exposure and a reduced shock avoidance of 15V, 30V, 45V and 60V (Fig. 1C). We did not find any effect for 90V shock avoidance indicating a limit for the modulation of shock avoidance. Moreover, this effect lasts for minutes to hours depending on the intensity of prior shock experience (Fig. 1D and fig. S2A). We then tested whether a prior shock experience also affects avoidance of innately aversive odors (Fig. 1E). We pre-exposed flies to different intensities of electric shock and then measured their capacities to avoid aversive odors. We found a linear correlation between shock pre-exposure and a reduced odor avoidance (Fig. 1E), confirming prior work (*33*). Furthermore, as for the effect on 30V shock avoidance, the effect of 60V pre-exposure on odor avoidance only lasted for less than 5 minutes (Fig. 1F). This indicates that 60V pre-exposure can alter both shock and odor avoidance with similar or different mechanisms. Importantly, the observed reductions of shock and odor avoidance following shock pre-exposure were not due to locomotor defects as shock pre-exposure of 90V or lower intensities did not alter flies’ climbing activity (fig. S2B). Overall, our findings demonstrate that a prior shock experience affects the innate value of future shocks and odors in an intensity-dependent manner, thereby altering odor-shock associative learning.

### Flies detect voltage intensity using *painless*, *piezo* and *straightjacket* genes

To understand how electric shock response can be modulated by a prior aversive experience, we started to delve into the sensory perception of electric shock. In both vertebrates and invertebrates, electric shocks have been used for decades as nociceptive stimuli to study nociception/pain (*29*, *34*, *35*), as stressful stimuli to study stress/anxiety/depression (*36*, *37*), and as negative reinforcement in the context of learning and memory (*37–39*). Although electric shock is ecologically questionable (*40*), but see (*41*), it has the advantage of being a quantifiable, reproducible and non-invasive nociceptive stimulus (*40*). However, the way flies perceive and integrate electric shocks remains puzzling. Flies respond to electric shock by freezing, jumping and increasing their locomotion to avoid shock (*38*, *42*), and this is dependent on voltage intensity (*19*). Shock delivery is therefore likely to activate mechanosensory receptor that could be, for instance, involved in muscular nociception. In addition, the resistance of the insect cuticle to electron displacement is likely really high between two (to six) parts (generally between legs) of the insect body (*43*). We found that flies’ shock avoidance is driven by the intensity of the voltage, not by the intensity of the current (fig. S2C). This indicates that it is the force pushing electron displacement and not the quantity of electrons moving in the fly body that is responsible for shock perception and avoidance. Thus, the higher the shock the more likely heat sensing will be involved.

Across phyla (*44*, *45*), TRP channel (i.e. *painless* in *Drosophila* (*46*)) and *straightjacket* (α_2_δ family calcium channel subunit; (*47*)) are necessary for both heat and mechanosensory perception, while *piezo* (*48*) is only involved for mechanosensory perception. We tested the role of these genes in shock avoidance, and using heat avoidance as a control. We found that the *straightjacket* and *piezo* mutants show significantly reduced shock avoidance independently of the intensity (fig. S3A). At 90V, heat is likely increased with relatively fixed cuticle resistance. Accordingly, *painless* is only necessary for high 90V shock avoidance (fig. S3A). As expected, only *painless* and *straightjacket* are necessary for heat avoidance (fig. S3B). These results indicate that an electric shock activates thermal and mechanosensory nociceptive pathways. Once integrated in the central nervous system, an electric shock triggers reflexive nocifensive behavior (e.g. jump; (*42*)). Then, relayed to the central brain via ascending circuits (*49*, *50*), electric shock induces shock avoidance and serves as a negative reinforcement signal during learning through the activation of dopaminergic neurons (*5*, *27*, *28*). In addition, shock triggers an acute stress response that may alter the perception and response of future nociceptive stimuli, as observed in vertebrates (*51*, *52*).

### Nociceptive stimulation triggers CO_2_ release inducing stress-dependent analgesia

How is electric shock nociceptive information modulated by prior experience? Can it be influenced by other stressful experiences? Can prior electric shock experience modulate response to nociceptive stimuli of a different modality? We addressed these questions and first found that heat avoidance was proportionally reduced following shock pre-exposure of different intensities (Fig. 2A). Similarly, a strong mechanical stimulation (shaking flies in a tube with a vortex) also significantly reduced ensuing shock and heat avoidance (Fig. 2B). Importantly, as observed for shock pre-exposure (fig. S2B), mechanical stimulation did not alter flies’ locomotor/climbing activity (fig. S4). These findings revealed a general nociceptive modulatory effect of prior stressful experiences on nocifensive (avoidance) response. In addition to stressful mechanical stimulation, electric shock is known to induce the production of social *Drosophila* stress odorant (dSO) (*31*). We therefore proposed that electric shock triggers dSO release that induces an acute stress response. Within a specific time window (Fig. 1D and fig. S2A), this stress state decreases the perception and response of ensuing nociceptive stimuli regardless of sensory modality, a phenomenon known as stress-induced analgesia (*8, 51–53*).

**Fig. 2.**
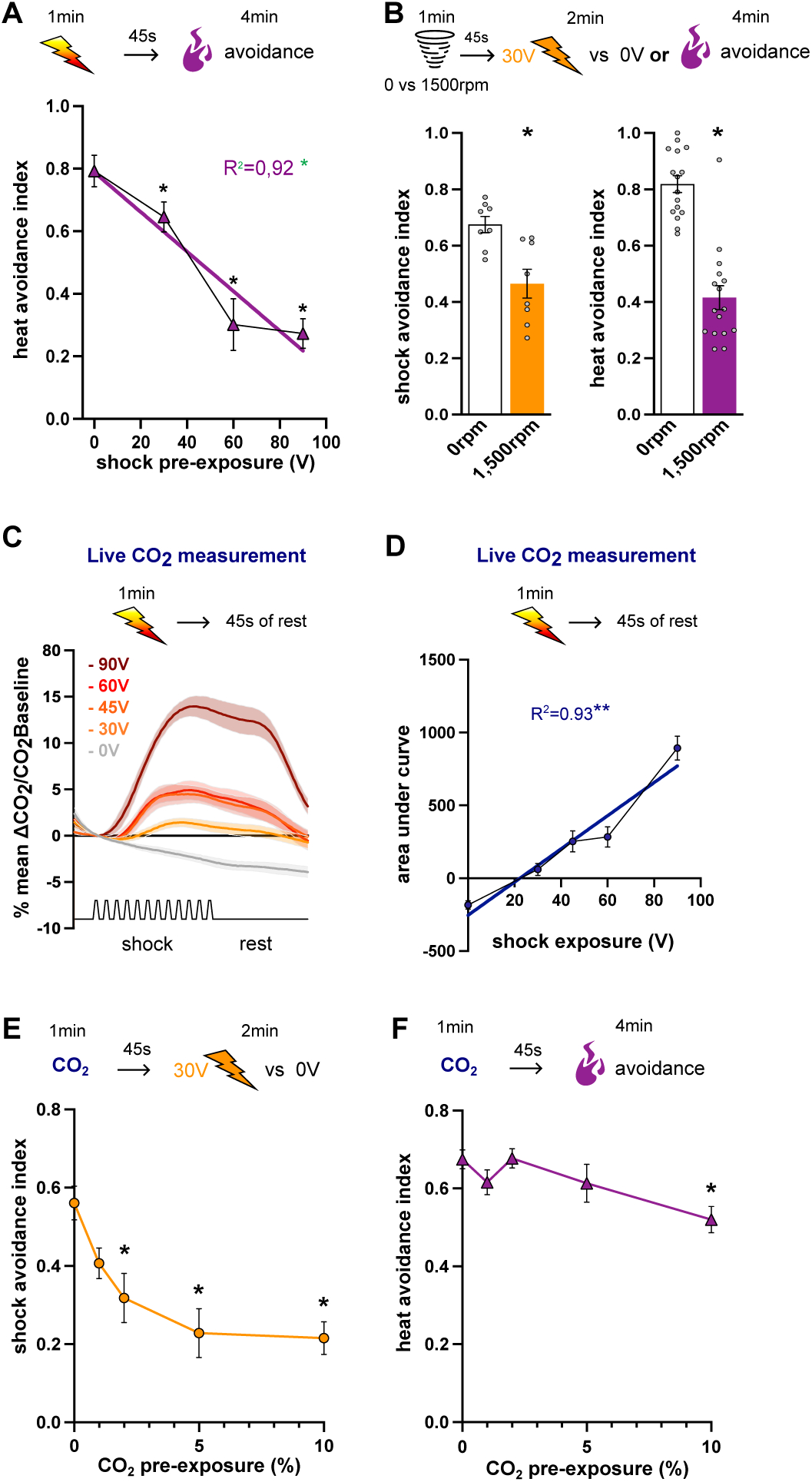
Heat and shock avoidance are decreased by shock, CO_2_ and mechanical stimulation. (**A**) Heat avoidance index (42 °C) linearly decreases after 30V, 60V or 90V shock pre-exposure, n=12. (**B**) Shock avoidance index (30V) significantly decreases after 1,500 rpm mechanical stimulation, n=8. Heat avoidance index (42 °C) significantly decreases after 1,500 rpm mechanical stimulation, n=12. (**C**) Mean percentage of CO_2_ production measured with a respirometer during 1 min of shock stimulation and 45 s of rest (**D**) Area under the curve of CO_2_ production in (C) is correlated with shock pre-exposure intensity, n=8-11. (**E**) Dose-dependent decrease of 30V shock avoidance index after 2%, 5% or 10% CO_2_ pre-exposure, n=13. (**F**) Heat avoidance (42°C) index significantly decreases after 10% CO_2_ pre-exposure, n=9-10. Top of each graph represents the protocol. Data presented as mean ± standard error of mean (SEM). Individual data points are displayed as dots. Asterisks indicate statistically significant differences (p < 0.05). Individual data points are displayed as dots. Asterisks indicate statistically significant differences (p < 0.05).

Mechanical stimulation can induce the production of dSO such as carbon dioxide (CO_2_) (*31*) (also see (*54*)), which flies either avoid or approach depending on the food source (*55, 56*) and the ecological context (*57*). We therefore reasoned that shocking flies might similarly trigger a dose-dependent production of CO_2_. Using a respirometer, we measured CO_2_ level from a group of flies exposed to different electric shocks intensities. CO_2_ production increased during and after electric shock stimulation, and is scaled with shock intensity (Fig. 2, C and D). These results were confirmed using gas chromatography and mass spectrometry (GC/MS) (*31, 58*) (fig. S5, A and B). Previous work showed that CO_2_ production can depend on fly density in confined environments (*58*) such as shock tubes. Consistent with this, pre-exposure to 60V shock reduced a subsequent 30V shock avoidance in high-density conditions (above 50 flies per 13.25 cm^3^ shock tube) (fig. S5C). This suggests interactions between shock intensity, fly density, CO_2_ accumulation and does-dependent analgesia. We tested this hypothesis by pre-exposing flies to different concentrations of CO_2_ and then testing shock avoidance. We found a dose-dependent effect with a significant reduction of 30V shock avoidance starting at 2% CO_2_ and reaching a plateau at around 5% (Fig. 2E). We noticed a discrepancy between the % CO_2_ scale used in behavioral experiments from that measured with the respirometer and GC/MS experiments (Fig. 2, C and D and fig. S5, A and B). This can be explained by three reasons as we sampled in the middle of the shock tube: 1) CO_2_ is 1.5 times heavier than air and could accumulate at the bottom of the tube, 2) when flies are shocked, they fall at the bottom of the tube where the CO_2_ concentration might be the highest, and 3) when flies are shocked they produce and accumulate CO_2_ in the tracheal system, while CO_2_ pre-exposure will only require flies to inhale CO_2_ mixed with air to reach similar concentration. Similarly, discrepancies between CO_2_ production vs behavioral effective concentrations have also been reported in Suh et al. (2004) (*31*). Remarkably, CO_2_ pre-exposure did not affect avoidance of a stronger 60V shock (fig. S5D), indicating limits of CO_2_-induced analgesia. Consistently, CO_2_ pre-exposure significantly reduced heat avoidance only at high (10%) concentration (Fig. 2F). Importantly, CO_2_ pre-exposure had no effect on climbing activity (fig. S5E), thus ruling out general locomotor impairment, which is only present when CO_2_ concentration is delivered above 60% (*59, 60*). Altogether, our results demonstrate that electric shock delivery induces intensity-dependent CO_2_ production which, perceived as a stress signal by themselves and conspecifics, induces dose-dependent analgesia. Our results also show that, as for shock-induced decrease of shock and heat avoidance (Figs. 1C and 2A) and the requirement of specific nociceptive channels (fig. S3), nociceptive modulation exhibits thresholds, likely arising from the combined contribution of sensory pathways, including the CO_2_-reponsive one.

### Olfactory CO_2_ perception is not involved in stress-induced analgesia

How is CO_2_ perceived and relayed to modulate nociception? Organisms can sense CO_2_ externally or internally via different mechanisms (*61–63*). Notably, flies can detect external CO_2_ via heteromeric receptor encoded by *Gr21a* and *Gr63a* genes expressed in a single population of antennal olfactory receptor neurons housed in basiconic ab1 sensilla (*31, 64, 65*). Atmospheric CO_2_ levels are usually around 0.04%, but at above 2%, CO_2_ can dissolve in antennal lymph fluid especially with H_2_O reducing local pH (CO_2_ + H_2_O <-> H_2_CO_3_ <-> HCO_3_^-^ + H^+^; (*66*)). So, above 2% CO_2_, flies use Gr21a/Gr63a (*64*) in combination with acid-sensing Ir64a-expressing neurons (*21*). We therefore tested the extent to which Gr21a/Gr63a-and Ir64a-olfactory sensing neurons (fig. S6, A and B) can relay CO_2_ stress signal emitted by flies receiving shocks to participate in nociceptive modulation. We expressed the inhibitory channel rhodopsin GtACR1 (*67*) in Gr63a-GAL4 and Ir64a-GAL4 and exposed flies to 525 nm green light during the 1 min of 60V pre-exposure as well as the following 45 sec of no shock, as CO_2_ is continuously emitted during and after shock stimulation (Fig. 2C). Our results showed that inhibiting Gr63a- or Ir64a-expressing neurons during pre-exposure had no effect on subsequebnt 30V shock avoidance (fig. S6 C and D). Therefore, olfactory CO_2_ Gr21a/Gr63a or acid Ir64a sensing circuits do not participate in CO_2_-induced analgesia.

### Td neurons are part of Gr28b.c-expressing neurons that target the brain and VNC

Flies must therefore quickly sense CO_2_ through dedicated circuits to directly modulate nocifensive behavior, here reflected by a reduced the avoidance of noxious stimuli. Previous work in larvae identified tracheal dendrite (td) neurons (*68, 69*) that respond to elevated CO_2_ (hypercapnia) and reduced O_2_ (hypoxia) within the respiratory tracheal network (*70, 71*). Respiratory gases enter and leave the tracheal system through closable valves called spiracles (*72*), located along the thorax and abdomen of the fly (fig. S7A). This gas exchange has discontinuous respiratory cycles alternating between periods of breath-holding, restricted oxygen uptake and burst of CO_2_ release. Although these cycles are largely driven by tracheal-to-environmental gas gradients, muscle contractions also contribute to it (for reviews see (*73–75*)). We hypothesize that when flies are shocked, they respond by jumping and increasing locomotor activity causing high metabolic CO_2_ production released through breathing (Fig. 2, C and D and fig S5, B and C). As CO_2_ production accumulates in the confined environment (shock tube) above the tracheal concentration, CO_2_ can then enter and accumulate in the tracheal system inducing hypercapnia. Critically, hypercapnia is a potent physiological stressor that has been shown to produce analgesia (*76, 77*), but the underlying mechanisms remain poorly known. We therefore hypothesize that flies sense stressful hypercapnia through td neurons, triggering nociceptive modulation (analgesia) resulted in reduced nocifensive shock and heat avoidance. However, td neurons have yet been described in adult flies.

We therefore characterized in detail the anatomy of Gr28b.c-GAL4, targeting td neurons in larvae (*71*). First, we expressed UAS-mCitrine (a strong membrane-bound mutated YFP) and UAS-DenMark::RFP (a dendritic marker; (*78*)) in Gr28b.c-GAL4. We reasoned that, as in larvae (*69, 71*), sensing CO_2_ may occur in the trachea near the spiracles. Screening through the cuticle at the thorax and the abdomen, we found dendrites overlapping with the trachea close to the thoracic spiracle sp2 (Fig. 3A and fig. S7A). Our 3D reconstruction clearly revealed dendrites in the trachea (Fig. 3B and fig. S4B), providing, to our knowledge, the first evidence of the presence of td neurons in adult flies. We then explored the innervation and projections of Gr28b.c-expressing neurons in the adult CNS. Gr28b.c-expressing neurons (containing td neurons) innervate the brain mostly in the suboesophageal zone (SEZ) with few antennal lobe interneurons (Fig. 3C). Gr28b.c-expressing neurons are also present in the ventral nerve cord (VNC) mostly located in the abdominal ganglia with some innervation in the center of the VNC and few neurons in the different neuromeres. Importantly, td neurons’ CNS projection in larvae are also visible in the SEZ and VNC, (*69, 70*). Other sensory neurons are targeted by Gr28b.c-GAL4 and project to the CNS (*79–81*) such as abdominal, proboscis and leg neurons (fig. S7, C-E). However, considering their function, these neurons are unlikely contributing to sensing acute volatile CO_2_. We therefore propose that td neurons are well placed to sense hypercapnia in the fly respiratory system and induce analgesia.

**Fig. 3.**
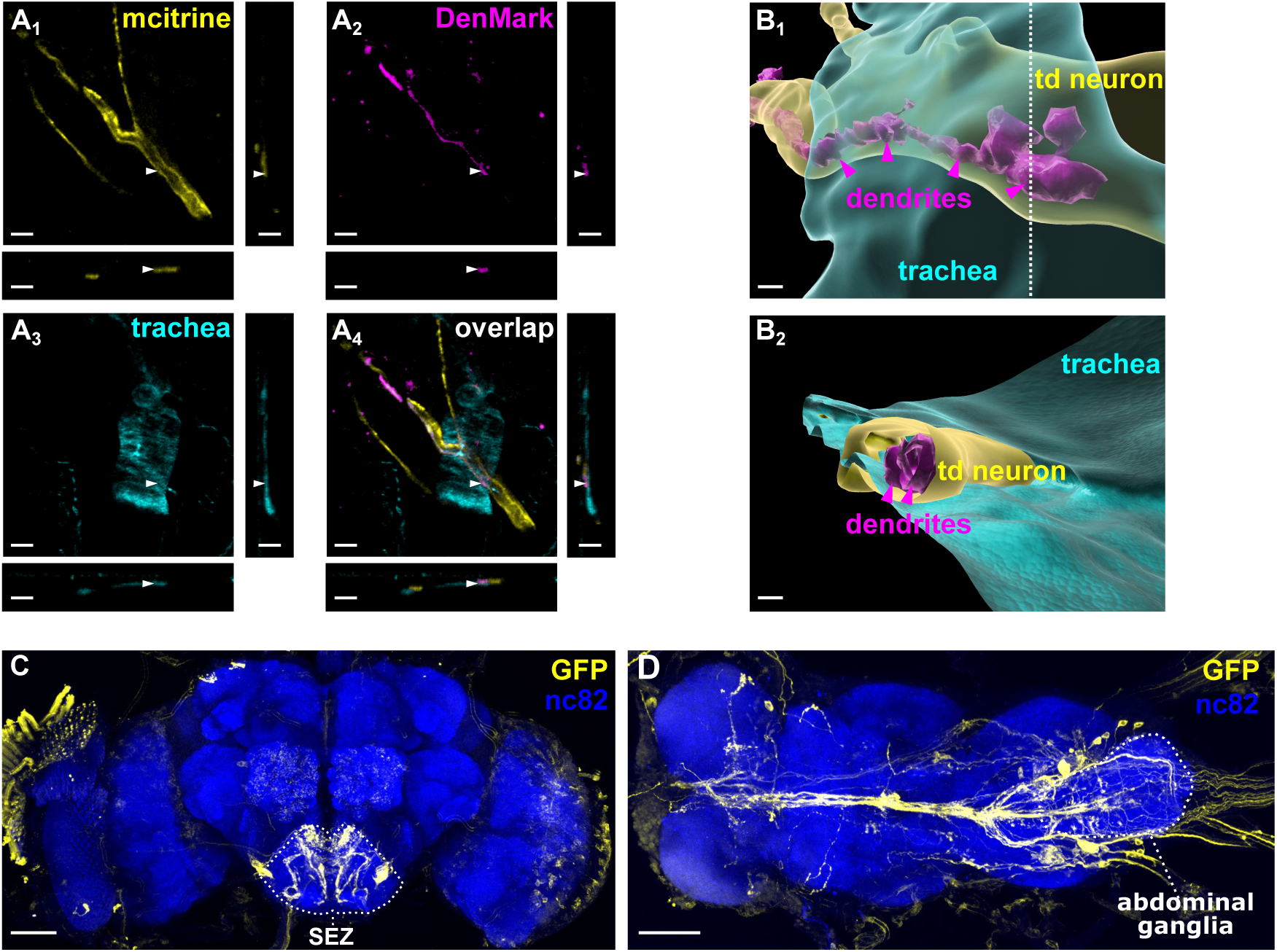
Gr28b.c-expressing neurons target tracheal dendrite (td) neurons. (**A**) Gr28b.c-GAL4 driving UAS-mCitrine (yellow) (**A_1_**) and UAS-DenMark::RFP (magenta) (**A_2_**) with a UV-responding trachea (cyan) (**A_3_**) and the overlap (**A_4_**). Scale bar is 10μm. (**B**) 3D reconstruction of Gr28b.c-GAL4 (td neurons) of images in (A) showing the location of dendrites along a neurite of td neurons (**B_1_**) in the trachea. (**B_2_**) sliced 3D reconstruction of images in B_1_ (at the white dot line) to expose dendrites in td neurons. Scale bar 3 μm for B_1_ and 5 μm for B_2_. (**C** and **D**) Gr28b.c-GAL4 driving UAS-mCD8::GFP (yellow) labelling td CO_2_ sensing neurons and other sensory neurons targeting the brain (**C**) and VNC (**D**). The whole brain and VNC is labelled with the presynaptic marker anti-Bruchpilot, nc82 (blue). Scale bars, 50 μm.

### Tracheal dendrite neurons relay hypercapnia to induce analgesia

To test the role of td neurons in relaying hypercapnia-induced analgesia, we expressed GtACR1 in Gr28b.c-GAL4 expressing neurons. We exposed flies to 525 nm green light during 10% CO_2_ pre-exposure as well as during the following 45 sec of no shock, and then performed 30V shock avoidance. Strikingly, optogenetic inhibition of Gr28b.c-expressing neurons during CO_2_ pre-exposure prevented the analgesic effect of CO_2_ on subsequent 30V shock avoidance (Fig. 4A). Importantly, inhibiting Gr28b.c-expressing neurons for the same duration during air pre-exposure did not alter 30V shock avoidance (Fig. 4B). We then tested the role of Gr28b.c-expressing neurons during shock pre-exposure on subsequent shock avoidance. Again, we found that inhibiting Gr28b.c-expressing neurons prevented the analgesic effect of 60V (Fig. 4C). Importantly, inhibiting Gr28b.c-expressing neurons for the same duration without shock pre-exposure did not alter 30V shock avoidance (Fig. 4D). Moreover, control experiments without 525 nm green light stimulation had no effect on 30V shock avoidance (fig. S8A). Gr28b.c-expressing td neurons therefore likely relay CO_2_ in the tracheal system to induce analgesia.

**Fig. 4.**
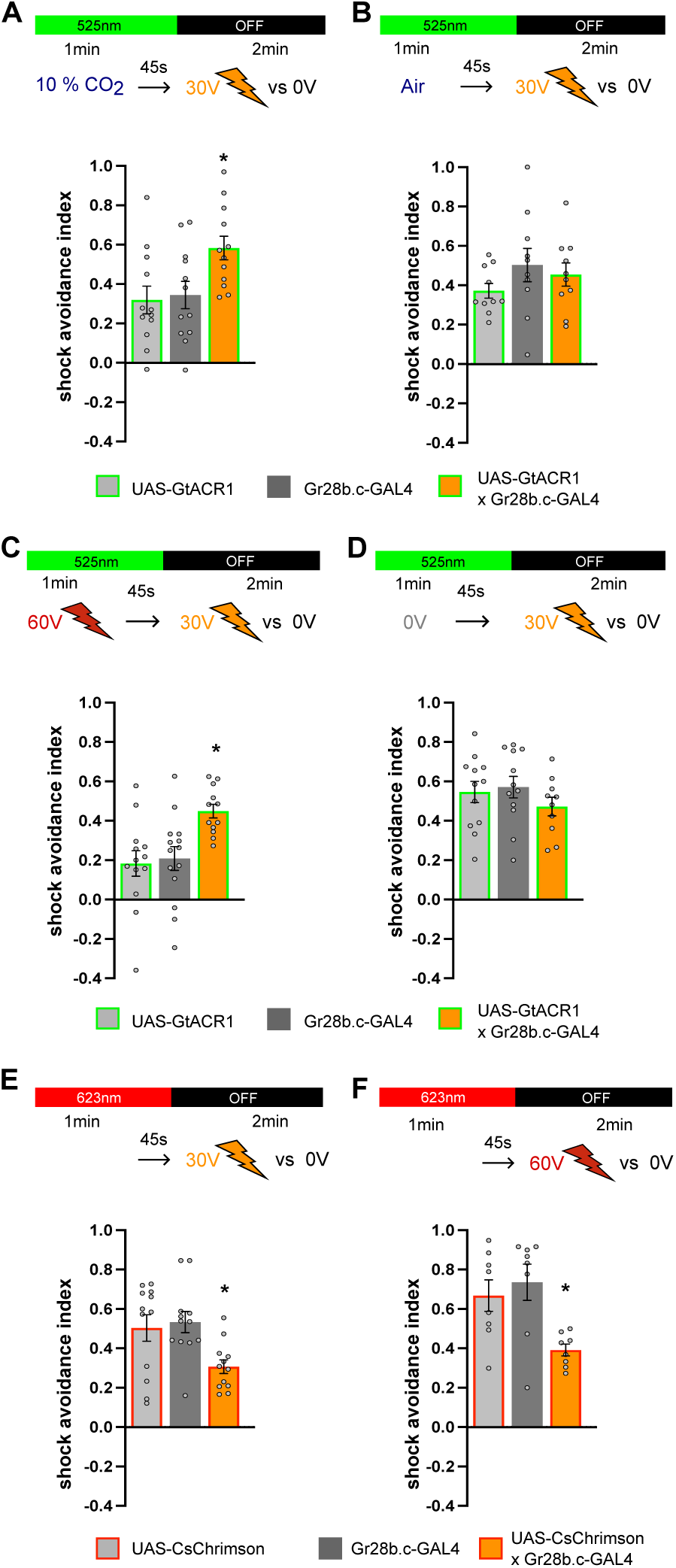
Gr28b.c-expressing (td) neurons are a key component of stress-induced analgesia. (**A**) Optogenetic silencing of Gr28b.c-expressing (td) neurons with GtACR1 during CO₂ pre-exposure (10%) restores normal 30V shock avoidance response, n = 12. (**B**) Optogenetic silencing of Gr28b.c-expressing (td) neurons with GtACR1 during air pre-exposure has no effect on 30V shock avoidance, n=10. (**C**) Optogenetic silencing of Gr28b.c-expressing (td) neurons with GtACR1 during 60V shock pre-exposure restore normal 30V shock avoidance, n=12-14. (**D**) Optogenetic silencing of Gr28b.c-expressing (td) neurons with GtACR1 during 0V shock pre-exposure has no effect on 30V shock avoidance, n=10-12. (**E**-**F**) Optogenetic activation of Gr28b.c-expressing (td) neurons with csChrimson during a 1 min pre-exposure significantly decrease following 30V shock avoidance, n=12 (**E**), but 60V shock avoidance, n=8 (**F**). Top of each graph represents the protocol. Data presented as mean ± standard error of mean (SEM). Individual data points are displayed as dots. Asterisks indicate statistically significant differences (p < 0.05).

Hypercapnia during shock/CO_2_ pre-exposure might activate td neurons to trigger nociceptive modulation and a change of nocifensive response. We tested this hypothesis by artificially activating Gr28b.c-expressing td neurons to mimic CO_2_ triggering stress-induced analgesia. We expressed UAS-csChrimson (*82*) in Gr28b.c-GAL4 and exposed flies to 633 nm red light during 1 min and 45 sec, and then performed 30V shock avoidance. Activating td neurons reproduced the effect of 60V shock/CO_2_ pre-exposure by significantly reducing 30V shock avoidance (Fig. 4E). Intriguingly, optogenetic activation of td neurons was sufficient to decrease subsequent 60V shock avoidance and heat avoidance (Fig. 4F and fig. S8B). As shown earlier, electric shocks of 45V and above (Fig. 1C), but not 10% CO_2_ pre-exposure (fig. S5D), alter 60V shock avoidance. We therefore propose that td neurons relay hypercapnia and potentially other signals (as a consequence of high shock stimulation) to the central nervous system to produce dose-dependent analgesia.

How do td neurons interact with neurons in the CNS to modulate nocifensive behavior? We propose that td neurons interact either with the nociceptive ascending pathway (nociceptor or ascending neurons) or with the descending avoidance pathway. To address these possibilities we characterized the postsynaptic partners of neurons labelled by Gr28b.c-GAL4 in the CNS and used the *trans*-Tango labelling strategy (fig. S9) (*83*). In the brain, we found signal in neurons innervating the SEZ and the superior medial protocerebrum (SMP) with some cell bodies in the pars intercerebralis (PI) (fig. S9A and movie S1). Interestingly, the SEZ output neuron 01 (SEZON01), known to relay nociceptive inputs to PPL1-γ1pedc punishment DANs in the SMP, has dendrites in the dorsal part of the SEZ (*84, 85*) where we observed our *trans*-Tango signal. In the VNC, we found most of the *trans*-Tango signal in the abdominal ganglia, including some ascending neurons (fig. S9B and movie 2). Importantly, nociceptors are known to project into the abdominal ganglia (*50, 86*) where ascending nociceptive neurons have their dendrites (*49, 50*). We also performed a GABA immunostaining on our *trans*-Tango signal and found that a subset of Gr28b.c downstream neurons in the brain and VNC are GABAergic (fig. S9, C-F). While Gr28b.c-expressing neurons are diverse (*79–81*), their CNS innervation suggests that CO_2_ activation of td neurons modulate nociceptive pathways, potentially via GABAergic inhibition. This modulation would ultimately reduce the representation of noxious signals the brain to *in fine* decrease avoidance behavior.

### CO_2_ pre-exposure decreases aversive value coding

Based on our findings, we propose that td neurons activated by hypercapnia decrease nociceptive information reaching the brain therefore decreasing punishment DANs response to noxious stimulation. In an intensity-dependent manner, this will then decrease aversive value assigned to experience during learning and ultimately reduce memory recall (Fig. 1, A and B). We used *in vivo* two-photon calcium imaging to test this hypothesis and recorded shock-evoked responses in PPL1-γ1pedc punishment DANs (known to relay shock response (*5, 27, 28*)) after CO_2_ pre-exposure. We expressed GCaMP6s in MB504B-GAL4 (*87*) targeting PPL1-γ1pedc and recorded 30V shock-evoked response after pre-exposing flies to air or 10% CO_2_ during 1 min (Fig. 5A). Consistent with our hypothesis, we found that 30V shock responses were significantly reduced by prior 10% CO_2_ pre-exposure (Fig. 5 B and C, and fig. S10A). Importantly, the same strategy did not alter neuronal response evoked by 60V shock stimulation (Fig. 5, D and E, and fig. S10B), in line with our behavioral results (fig. S5D). Altogether, these findings show that CO_2_ pre-exposure selectively attenuates nociceptive signaling in PPL1-γ1pedc punishment DANs, likely reducing aversive value coding in the MB.

**Fig. 5.**
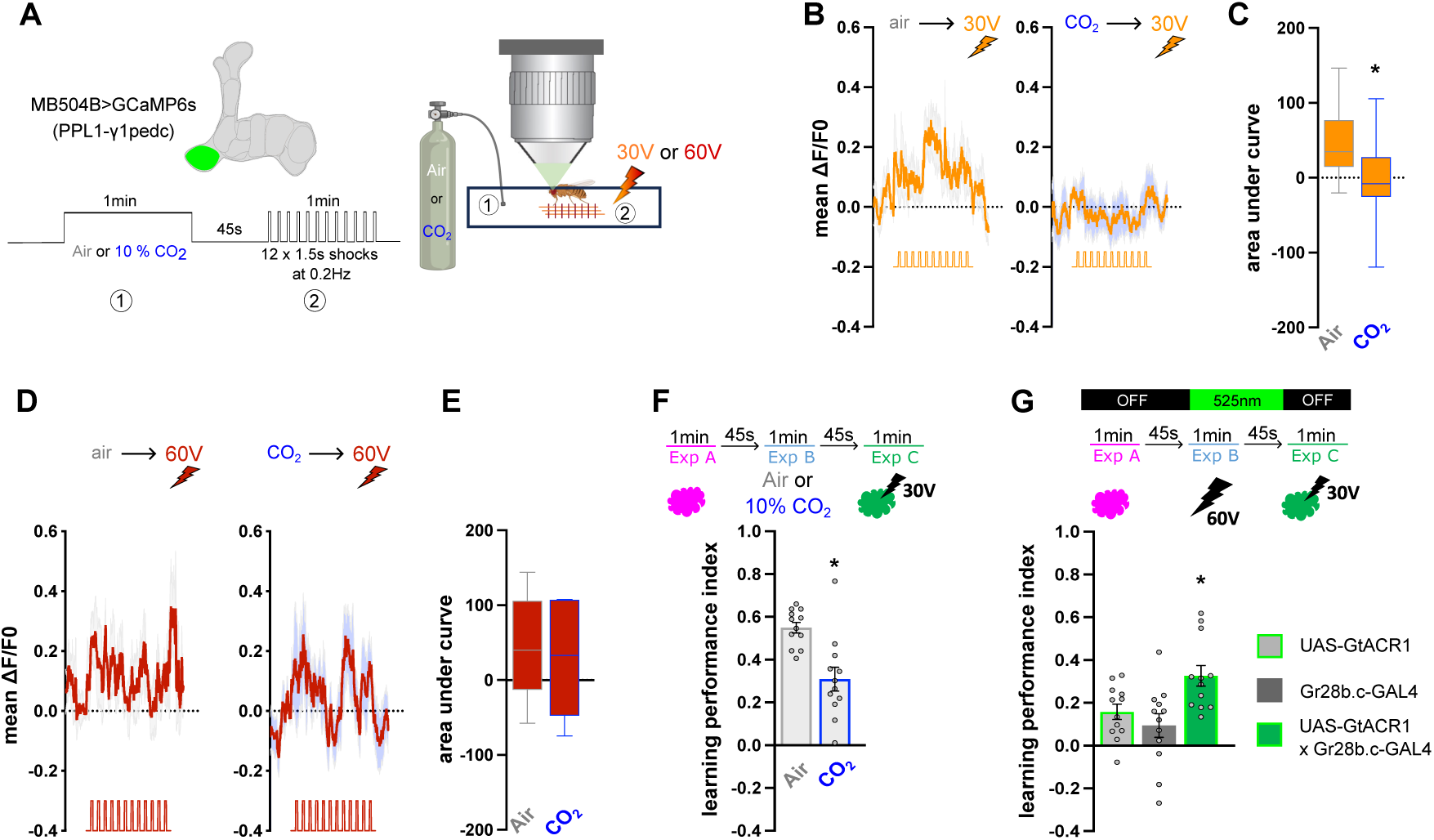
CO_2_ pre-exposure decreases aversive value coding. (**A**) Experimental setup for *in vivo* calcium imaging. Flies are presented with electric shocks to the legs, while neural activity is recorded in punishment PPL1-γ1pedc DANs targeted by MB504B-GAL4 expressing GCaMP6s. (**B**) Mean ΔF/F0 calcium transients ± standard error of the mean (SEM) measured from PPL1-g1pedc to 30 V shocks after air or 10% CO_2_ pre-exposure, n=11. (**C**) Mean area under curve calcium transient from PPL1-γ1pedc response to 30 V shocks is significantly decreased after 10% CO_2_ pre-exposure, n=11. (**D**) Mean γF/F0 calcium transients ± standard error of the mean (SEM) measured from PPL1-γ1pedc to 60 V shocks after air or 10% CO_2_ pre-exposure, n=7-8. (**E**) Mean area under curve calcium transient from PPL1-γ1pedc response to 60 V shocks is not affected by 10% CO_2_ pre-exposure, n=7-8. (**F**) Pre-exposing flies to 10% CO_2_ prior to associating odor C + 30V decreases avoidance of learned odor C during the memory test, n=12. (**G**) Optogenetic inhibition of Gr28b.c-expressing (td) neurons driving GtACR1 during 60V pre-exposure rescue the 30V aversive value assigned to odor C during learning, n=12. Top of each graph represents the protocol. Individual data points are displayed as dots. Data presented as mean ± standard error of mean (SEM). Data in (**C**) and (**E**) are presented as boxplot with the min, median and max value. Asterisks indicate statistically significant differences (p < 0.05).

Together, these data therefore explain our results in Fig. 1B, in which odor C is assigned a lower negative value when preceded by a strong 60V shock stimulation that triggers important CO_2_ release (Fig. 2, C and D). Accordingly, pre-exposing flies to 10% CO_2_ prior to odor C + 30V association similarly reduced learned avoidance of odor C during the memory test (Fig. 5F). Finally, to directly test our model, we found that optogenetic inhibition of td neurons during the 60V pre-exposure restores the aversive value assigned to odor C during learning (Fig. 5G). Importantly, control optogenetic inhibition of td neurons during a 0V pre-exposure control condition had no effect on 30V aversive value assigned to odor C during learning (fig. S11A). Moreover, control flies not exposed to light stimulation still exhibited normal reduction in aversive learning following 60V pre-exposure (fig. S11B).

Overall, we propose a model (Fig. 6) in which noxious stimulation triggers CO_2_ production that accumulates in a confined environment and in the tracheal system to induce hypercapnia. Detected by td neurons, hypercapnia in turn reduces nociceptive inputs reaching the brain and therefore decreases nocifensive avoidance response. Hypercapnia-induced analgesia therefore decreases nociceptive input to punishment DANs assigning a lower aversive value to olfactory stimuli during learning, ultimately reducing learned avoidance response.

**Fig. 6.**
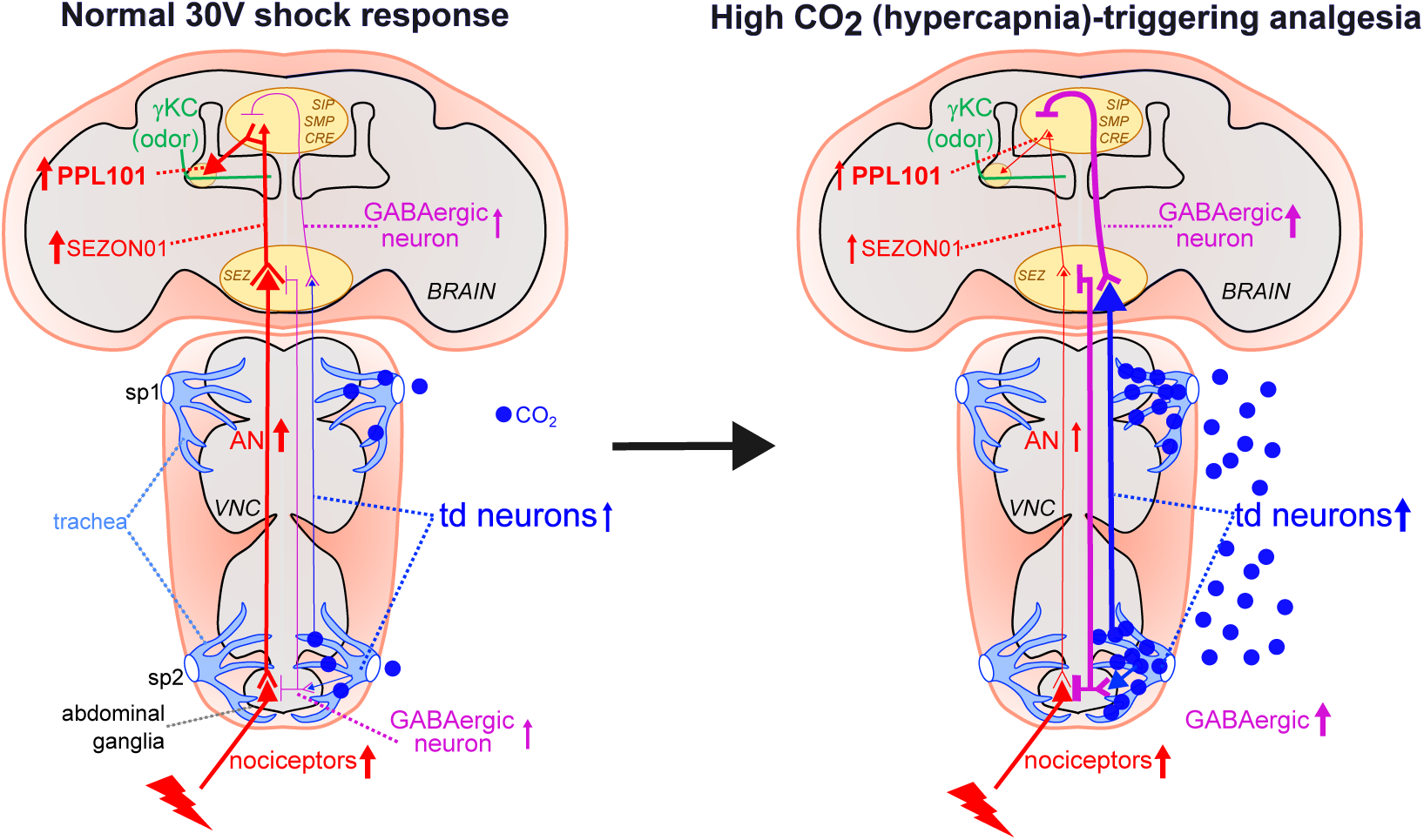
Circuit model of hypercapnic stress modulation of aversive value coding. High CO_2_ (hypercapnia) is detected by tracheal dendrite (td) neurons innervating the ventral nerve cord (VNC) (potentially the abdominal ganglia) and/or suboesophageal zone (SEZ) in the brain. Td neurons activate GABAergic neurons innervating the VNC and/or SEZ. These GABAergic neurons (activated by hypercapnia) interact with nociceptors and ascending nociceptive neurons in VNC and/or superior medial protocerebrum (SMP) or in the SEZ with ascending nociceptive neurons or SEZ output neuron 01 (SEZON01; relaying nociceptive information). Hypercapnia relayed by td neurons therefore decreases nociceptive information reaching punishment dopaminergic neurons (e.g. PPL01/PPL1-γ1pedc) therefore decreasing aversive value coding.

## DISCUSSION

Our study reveals that tracheal dendrite (td) neurons, detecting hypercapnic stress in the tracheal network, are a key modulator of nociceptive information thereby altering the way flies learn and respond to dangerous stimuli. We first show that a prior noxious/stressful experience induces analgesia. Then we show that noxious stimulation induces the production of CO_2_, likely via hypermetabolic production and its release through breathing. Accumulation of CO_2_ in the confined surrounding environment and in the respiratory system causes hypercapnia detected by Gr28b.c-expressing td neurons. Potential interaction with nociceptor/ascending pathways in the CNS decreases brain representation of threatful stimuli, therefore inducing analgesia measured by a significant decrease of noxious stimulus response. Modulation of d neurons also changes the way punishment DANs respond to noxious stimuli therefore altering aversive value coding and the formation of aversive memory. This work provides the first circuit model of stress-induced analgesia in *Drosophila*, improving our understanding of how a prior stressful/hypercapnic experience can modulate nociceptive information in the CNS and aversive value coding in the brain.

### Td neurons, a component of the fly vagus nerve detecting noxious signals in the respiratory system?

In mammals, the lower airways (trachea, bronchi and lungs) receive more than 90% of their sensory innervation from the vagus nerve (*88–90*). Most of those afferent nerves are unmyelinated, slowly-conducting C fibers which are activated by noxious or potentially noxious stimuli (*91*). Intriguingly, vagus nerve stimulation can change pain perception (*92, 93*). In adult flies and larvae, a vagus nerve has been described connecting the enteric system and the brain (for reviews see (*94–96*)). Our work proposes that fly td neurons are a component of a fly vagus nerve that detect high interoceptive CO_2_ inducing a stress state (hypercapnia) leading to increased breathing (releasing more CO_2_) and triggering analgesia by interaction with the nociceptive pathways. In larvae, CO_2_ can be then detected by Gr28b.c expressed in td neurons (*71*). In the trachea, CO_2_ is likely mixed with water producing H+, HCO_3_^-^ and carbonated water. Td neurons could therefore indirectly detect CO_2_ as acid or bicarbonate via specific receptors which are not Gr21a/Gr63a or IR64a (fig. S6), but could also detect carbonation via Ir25a, Ir56d and Ir76b (*97, 98*). Hypercapnia often occurs alongside hypoxia (low oxygen levels). However, the level of CO_2_ detected (Fig. 2, C and D, and fig. S5B) or used in our pre-exposure experiments (Fig. 2E) are unlikely to affect oxygen level enough to produce hypoxia. As a vagus nerve component, td neurons could therefore detect noxious interoceptive signals via different receptors to trigger adaptive behavioral responses and analgesia, similar to what occurs in mammals (*92, 93, 99, 100*).

### SEZ vs abdominal ganglia nociceptive modulation

Building on prior work in larvae from Lu et al., (2024) we described the expression pattern of the Gr28b.c-expressing neurons in the periphery (at the tracheal level near the spiracles) and CNS of the adult fly. Although we have not fully mapped the connection between dendrites within the tracheal system and their axonal targets in the CNS, we propose that td neurons target both the SEZ and VNC, consistent with work in larvae (*69, 70*). Our *trans*-Tango experiment suggests that td neurons innervating the SEZ and VNC target directly or indirectly the nociceptive pathways: nociceptors, ascending nociceptive neurons or first relay in the SEZ (SEZON01) to control nociception (Fig. 3 and 6). In larvae, two distinct td neurons subsets target either the SEZ or VNC (*69, 70*). It is therefore likely that adult flies also possess at least two different subsets of td neurons targeting SEZ or VNC to modulate nociception and nocifensive response. In prior work, Gr28b.c-GAL4 was shown to be expressed in different organs of the fly body (*79–81*). However, although we used the GAL4 deposited by John Carlson, we do not find similar expression pattern in the brain and VNC (*80, 81*) (fig. S7, C-E). This discrepancy could be explained by difference in expression resulting from genomic insertion site (second or third chromosome). We also did not find similar expression pattern of Gr28b.c-expressing neurons in the brain as in Amrein and Thorne (2008) which might be explained by the use of a different GAL4 line. While Gr28b.c-GAL4 expression in other organs could account for part of the expression we observed, its presence in multidendritic neurons, abdominal wall, reproductive organs, Johnston organ, aristae and leg taste neurons are unlikely contributing to sensing volatile CO_2_.

Ascending neurons could express tachykinin receptors in the abdominal ganglia (*101, 102*), with substance P being a highly conserved member of the tachykinin neuropeptide family released by nociceptors to the first relay in the CNS (*103, 104*). In larvae, td neurons targeting interneurons then project onto neuroendocrine cells such as Diuretic Hormone 44 (DH44) and Corazonin (CRZ) neurons, both of which respond to 20% CO_2_ exposure (*70*). Intriguingly, DH44 (a homologue of mammalian corticotropin-releasing hormone) and CRZ (a homolog of mammalian Gonadotrophin Releasing Hormone) are known to regulate stress responses (*105, 106*). Similar connectivity could exist in adult flies involving neuroendocrine (DH44/CRZ) neurons in regulating CO_2_-induced modulation of nociceptive inputs and other functions. Furthermore, it remains unknown if other td neurons in adult flies that do not express Gr28b.c might also be involved in modulating nociceptive information. We also do not know whether the same or different td neurons innervate SEZ and VNC (Fig. 6).

### Stress and sensory modulation

In response to noxious stimulation (e.g. shock) flies jump and increase locomotor activity (*42*), which results in the metabolic production and accumulation of CO_2_ in the tracheal system and releasing it through breathing. Flies also produce and emit a blend of other dSO (*31, 54*). While flies usually avoid CO_2_ (but see (*55–57*)), they avoid it even more strongly when it is combined with other dSO (*31*). However, the presence of CO_2_ is essential for dSO avoidance, as flies do not respond to dSO if their ability to detect CO_2_ is inhibited (*56*). The olfactory CO_2_ pathway therefore likely alters the way other non-dSO are encoded and processed in the brain, notably in the lateral horn, involved in innate olfactory behavior (*107*). We found that CO_2_ concentration increases with shock intensities (Fig. 2, C and D, and fig. S5B). The decreased odor avoidance after shock stimulation (*33*) (Fig. 1E) could be due to a dose-dependent effect of CO_2_ (and potentially other dSO) increasing activity in the lateral horn (*108*). A general state of acute stress can also reshape responses to sensory stimuli (*109, 110*). The activation of the hypercapnia-induced neuroendocrine system in response to acute stress could also participate (with or without the olfactory CO_2_ pathway) in the modulation of odor processing and behavioral responses.

### Mechanisms for stress-induced analgesia

The effects of stress on pain are bidirectional and depend on its nature, duration, and intensity, with stress capable of either attenuating pain (stress-induced analgesia) or intensifying it (stress-induced hyperalgesia). Notably, the same stressor can induce analgesia when acute and, when uncontrolled and chronic, can become a potent trigger of pain and anxiety disorders leading to impairment of learning and decision-making processes (*51, 111, 112*). Here, we showed a CO_2_ pre-exposure effect on low-intensity (Fig. 2E) but not high-intensity (fig. S5D) shock avoidance. However, we showed that high-intensity shock stimulation (Fig. 1C) or artificial activation of td neuron (Fig. 4F and fig. S8B) exerted more pronounced effects on responses to high 60V shock and heat. Accordingly, we suggest that strong shock stimulation activates not only CO_2_-mediated hypercapnic stress pathways but also additional mechanisms, such as diffuse noxious inhibitory controls (DNIC) (*113*). Because high-intensity shocks recruit both mechanosensory and thermosensory systems (fig. S3), they could trigger DNIC, which utilizes distinct complementary (e.g. descending) circuits and mechanisms of nociceptive modulation compared to the hypercapnic activation of td neurons.

### Stress and memory

Stress can profoundly shape memory processes. Notably, rapid neuroendocrine responses to stress can act from minutes to hours, thereby affecting function and plasticity within the mammalian HPA axis, prefrontal cortex, amygdala, striatum and ventral tegmental area (*114–116*). Interestingly, the amygdala can directly sense hypercapnia and acidosis, altering amygdalar neuronal (and likely glial) activity and inducing changes in fear-related behavior (*117*) and potentially the fear engram itself (*118*). Neuropeptides, like NPY, can also be stress-triggered and act in different brain structures to alter behavior (*119*). Therefore, by driving large-scale interactions between brain regions, stress can alter not only how information is encoded (as demonstrated here and discussed above) but also how it is stored and retrieved (*13, 120*).

In flies, the neuroendocrine and neuropeptide systems target many brain neuropils in response to stressors (*104, 121*). These signals could rapidly affect nociceptive sensory processing and learning-induced plasticity in the MB network as well as neural activity in the central complex (*35*) thereby modulating the expression of learned behavior. We previously demonstrated that the reward dopaminergic system innervating the MB is required to compare aversive experiences and encode a relative “better than” aversive value (*5*). Our current findings expand upon this suggesting that learning relative aversive value also depends on stress-related vagus nerve modulation and the neuroendocrine and neuropeptide systems.

## MATERIALS AND METHODS

### Fly strains and maintenance

Fly stocks were cultured at 21°C on standard cornmeal food which was made with 40L of tap water, 280g of agar, 1.5kg of yeast, 2kg of corn flour, 2.8kg of Dextrose, 120g of Tegosept and 460mL of Ethanol. Genotypes and sources of the fly lines used in this study are denoted in the **Table S1**.

### Behavioral experiments

For behavioral T-maze experiments, groups of ∼100 (or otherwise mentioned) 7-10 days old flies were housed for 18-24 h before training in a 25 ml vial containing standard cornmeal food and a 20 x 60 mm piece of filter paper at 23°C and 65% relative humidity and a 12h/12h light-dark cycle (except for optogenetic experiments for which flies were raised in the dark prior to the experiment).

For csChrimson experiments, 5-7 days old flies were placed in the dark on a 1mM all trans-Retinal solution mixed with fly food for 2 days prior to experiments. For heat avoidance assay, only 40 flies were selected and housed 18-24 h before in a 25 ml vial.

Electric shocks stimulation consisted of 12 pulses of 1.5 s delivered at 0.2 Hz. Various voltage intensities were delivered with a Grass S48 Square Pulse Stimulator (Grass Technology). Various current intensities were delivered with a combination of DS3 (90V current stimulator) and DG2A (trained/delay generator) (Digitimer Ltd, UK).

Conditioning odors were 3-octanol (OCT: 8 μl in 10 ml mineral oil), 4-methylcyclohexanol (MCH: 10 μl in 10 ml mineral oil), and isoamyl acetate exclusively as odor B (IAA: 16 μl in 10 ml mineral oil), with only 3-octanol and 4-methylcyclohexanol used during memory testing.

### Aversive olfactory conditioning

Aversive olfactory conditioning was performed in a T-maze using a training tube (of 13.25 cm^3^) equipped with an internal copper grid, as previously described (*5*). Conditioning was performed in three phases: In phase A, flies were exposed to odor A for 1 min without electric shock, followed by 45 s of clean air. In phase B, flies were exposed to one of the following conditions: exposure to odor B unpaired, to odor B paired with electric shocks (30V or 60V), exposure to electric shocks alone (30V or 60V), exposure to compressed air, or exposure to 10% CO_2_. In other experiments, targeted neurons were optogenetically inhibited using GtACR1. In phase C, flies were finally exposed to odor C for 1 min paired with 30 V electric shocks. Memory tests were performed immediately after training and were assessed in the dark using a 2 min two-odor choice (A vs C) in the T-maze. A Performance Index (PI) was calculated as the number of flies making the correct choice minus the number of flies making the wrong choice, divided by the total number of flies in each experiment. A single PI value is the average score from flies of the identical genotype tested with the reciprocal reinforced/non-reinforced odor combination.

### Pre-exposure paradigms

Pre-exposure consisted of exposing flies in a tube (of 13.25 cm^3^) to one of the following stimuli during 1 min (or otherwise mentioned) prior to testing shock/heat/odor avoidance: no stimulation (0V), or 15V, 30V, 45V, 60V, 90V electric shocks, compressed air or controlled CO_2_ concentrations ranging from 0% to 10% diluted in compressed air (controlled using two Mass Flow Controller, ANALYT-MTC GMBH, Germany), aversive odors, mechanical vortex stimulation (at 1,500 rpm, Fisherbrand ZX3 vortex mixer), optogenetic activation or inhibition of targeted neurons. Unless otherwise specified, the interval between pre-exposure and shock/odor/heat avoidance was 45 s.

### Shock avoidance assay

Shock avoidance was assessed in the T-maze by allowing flies to choose for 2 min in the dark between a shock tube delivering 24 electric shocks (at 0.2 Hz) and a shock tube without shocks. Shocks were delivered alternatively on the left or right shock tube to control for side bias. A shock avoidance index was calculated as the number of flies making the correct choice minus the number of flies making the wrong choice, divided by the total number of flies in each experiment.

### Odor avoidance assay

Odor avoidance was assessed in the T-maze by allowing flies to choose for 2 min between a tube containing an odor at a defined concentration and a tube containing solvent (mineral oil: MO). Odor-specific pre-exposure effects were tested using 4-methylcyclohexanol and acetophenone at concentrations ranging from 0 to 10^-3^. Odors were delivered alternatively on the left or right test tube to control for side bias. Odor avoidance index was calculated as the number of flies making the correct choice minus the number of flies making the wrong choice, divided by the total number of flies in each experiment.

### Heat avoidance assay

Heat avoidance was performed as previously published (*47*). The system consists of a transparent circular behavioral chamber (1.0 cm height, 7.0 cm diameter) and a variable heat element (model Isotemp RT Dgti HP230V, Fisherbrand). A temperature gradient is created in the chamber, from 42°C at the bottom to 31°C at the top. All tests were performed under low red light. 40 flies were placed into the behavioral chamber for 4 min of the heat avoidance test. Immobilized flies at the bottom of the chamber were counted as “incapacitated”. A heat avoidance index was calculated as the number of flies avoiding heat minus the number of flies incapacitated, divided by the total number of flies in each experiment.

### Optogenetic manipulations

For optogenetic experiments, a transparent shock grid was fabricated from a 75 mm × 45 mm PET sheet coated with a 175 nm ITO film (80 Ω/sq; Diamond Coatings Ltd., UK). Lanes 0.5 mm wide, separated by 0.3 mm gaps, were laser-etched to isolate electrodes. GtACR1 stimulation was delivered with four green LEDs (λ = 525 nm, 101.85 mW/mm²; Prolight Opto, PM2B-3LxE-SD, Taiwan) continuously illuminating the tube during shock/CO_2_ presentation. csChrimson stimulation was provided by four red LEDs (λ = 623 nm, 229.18 mW/mm²; Prolight Opto, PM2B-3LRE-SD, Taiwan) at 0.5 Hz, 1 s pulses illuminating the tube.

### Climbing assay

10 flies were put into a 25mL empty food tube, gently tapped on a mat on the bench (all flies falling at the bottom), and left 10 s to enable them to reach a red line drawn on the tube at 2.5 cm from the bottom. A percentage of flies reaching the red line in 10 s is then calculated.

### CO_2_ measurement

Live emission of CO_2_ was measured using LI-850 CO_2_ analyzer/respirometer. Groups of 100 flies were placed in a shock tube, which was flushed with a constant air flow at 80 mL/min. The amount of CO_2_ is measured during 1 min of shock stimulation and 45s of rest. For subsequent analyses custom MATLAB (MathWorks, Inc) scripts were used to first calculate variation of CO_2_ from the baseline (called CO_2_Baseline) with the following equation: ΔCO_2_ = ((CO_2_(t) – CO_2_Baseline) / CO_2_Baseline) x 100). The baseline was defined as the mean of CO_2_ produced during 2s before the start of the electric shock sequence. We then calculated the area under the curve (integral ΔCO_2_) during 1 min of shock stimulation and 45s of rest.

For gas chromatography/mass spectrometry, the analyses were performed on a GC Trace 1300 (Thermo, SN: 71800174), equipped with a GS-GASPRO column (Agilent) 60 m x 0.32 mm i.d. The spectrometer coupled to the chromatographic chain is an ISQ7000 (Thermo, SN: N2012006), equipped with a 70eV EI source. The mass spectrum was recorded between 10 and 300 Da. The CO_2_ signal was monitored by recording the SIM signal of the 44 Da ion. Flies were shocked for 1 min, then 20 µl of gas sample from the middle of the shock tube was introduced directly into the SSL injection module using a Hamilton Gastight syringe (50 µl #1705). Injections were performed in 1/100 split mode at a temperature of 200°C. The carrier gas was helium at 1.4 ml/min. Separation was performed with an isotherm at 50°C for 4 minutes. For each group, a measurement of the tube containing the flies before shock stimulation was taken and used to normalize the measurement after shock stimulation. CO_2_ of 1, 2, 5 and 10% (Chemlys, France) were used for the calibration.

### Immunohistochemistry and confocal microscopy

#### Brain and VNC

7-10 days old fly brains and VNC were dissected in ice-cold Schneider’s insect medium and fixed in 2% paraformaldehyde solution (Polysciences diluted in Schneider’s insect medium) at room temperature for 55 min. Fixed brains were then washed 3 x 20 min in PBS-triton (PBST) 0.5% and incubated in a blocking solution in 5% goat serum (Gibco^TM^) in PBS-triton (PBST) 0.5% for 90 min at room temperature. Brains were then incubated with primary antibodies, α-Bruchpilot, nc82, (1:30, mouse, DSHB) and chicken anti-GFP (1:1000) in PBST for 72h at 4°C. After 3 x 10 min washes in PBST, brains were incubated with secondary antibodies, Alexa Fluor 647-conjugated goat α-mouse 48h at 4°C. After 3 x 10 min washes in PBS, brains were mounted in Vectashield (Vector Labs) on a glass slide.

#### Whole fly mounting for visualizing tracheal dendrite neurons

7-10 days old flies (Gr28b.c expressing mCitrine and DenMark::RFP) were anesthetized on ice, mounted in Fluoromount-GTM polymerizing mounting medium on a glass slide. Tracheal signal is coming from autofluorescence using a UV laser stimulation.

#### For the trans-Tango experiment

7-10 old female flies, maintained at 25°C, were dissected in cold PBS and fixed for 55 min in 2% paraformaldehyde (Polysciences, diluted in PBS) rotating at room temperature. All subsequent incubation and washing steps were done while rotating, in the dark. Brains and VNCs were washed 3 x 10 in PBST 0.5% and incubated in blocking solution in 5% normal goat serum in PBST 0.5% for 60 min at room temperature. Brains and VNCs were incubated in primary antibody mixture (chicken anti-GFP 1:1000; rabbit anti-DsRed 1:500; mouse anti-nc82 1:25) and for the GABA-staining (chicken anti-GFP 1:1000; rat anti-HA 1:500; rabbit anti-GABA 1:500) for 48 h at 4°C. After 3 x 10 min washes in PBST, samples were incubated in secondary antibody mixture (goat anti-chicken Alexa Fluor 488; donkey anti-rabbit Alexa Fluor 568; donkey anti-mouse Alexa Fluor 647, all at 1:200) and for the GABA-staining (goat anti-chicken Alexa Fluor 488; goat anti-rat Alexa Fluor 555; donkey anti-rabbit Alexa Fluor 647, all at 1:200) for 48 h at 4°C. After 4 x 10 min washes in PBST, brains and VNCs were mounted in VectaShield (Vector Labs) on a glass slide.

Brains and VNCs were imaged on a Leica SP8 confocal microscope with a 20×objective and a 40×for close ups. Image resolution was 2048 x 1024 and 1024 x 512 and 1024 x 1024 for close ups with 1 µm step size and a frame average of 2. All images were analyzed using Fiji (*122*) and IMARIS software (Bitplane AG, Zurich, Switzerland).

#### *In vivo* two-photon calcium imaging

Functional-imaging experiments were performed as described previously (*5*) with some minor modifications. 1-2-day old flies were transferred into vials containing standard food (maximum of 30 flies per vial) and imaged 7-10 days later. Flies were briefly immobilized on ice (1-3 min) and mounted in a custom-made chamber allowing free antennae and leg movement. The head capsule was opened under room temperature buffer solution (5 mM TES, 103 mM NaCl, 3 mM KCl, 1.5 mM CaCl_2_, 4 mM MgCl_2_, 26 mM NaHCO_3_, 1 mM NaH_2_PO_4_, 8 mM Trehalose, 10mM glucose, pH 7). Membranes and trachea above the recording areas were manually removed. Individual flies in the recording chamber were placed under the 20x objective of a two-photon microscope (Zeiss LSM 710mp) and an electric shock grid (in copper) was positioned in contact with the fly’s legs (visualized with a camera AV MAKO U-029B (Stemmer Imaging)). For all flies, fluorescence signal was measured in a randomly chosen brain hemisphere. Fluorescence was excited by a Ti-Sapphire laser (Chameleon Ultra II, Coherent) using ∼140fs pulses, 80MHz repetition rate and centered at 920 nm. Images of 256 x 256 pixels were acquired at 6.34 Hz controlled with Zen software (Zeiss). CO_2_ stimulation was delivered from a 10% CO_2_ (containing 20.90% O_2_ and 69.3% N_2_) bottle connected to the fly chamber with a tubing at 3 psi. Electric shocks were delivered with a DS2A isolated voltage stimulator controlled with a DG2A train/delay generator (both from Digitimer, Ltd). Imaging (Zen software) and electric shocks were controlled via TTL signals using an Arduino board (Arduino uno Rev3, Arduino.cc) combined with a project board (K&H-102) and Arduino custom-made codes.

For CO_2_ effect on punishment DANs electric shock responses, each fly was recorded for 30 s before the onset of a 1 min of air or 10% CO_2_ pre-exposure followed by 45 s of fresh air and then a 1 min sequence 12 x 1.5 s electric shocks of a given voltage at 0.2 Hz.

Recorded images were manually segmented with Fiji using a custom-made code including an image stabilizer plugin (*123*). For each recording, one region of interest (ROI) was drawn around the zone expressing GCaMP6s axons of PPL1-γ1ped targeted by MB504B after image stabilization to generate the summed fluorescence at each frame. A second ROI of the same size was chosen in the background where no changes occur during the whole recording. The GCaMP6s fluorescence (F(t)) was then calculated by subtracting the background (mean fluorescence of the second ROI). For subsequent analyses custom MATLAB (MathWorks, Inc) scripts were used to first calculate variation of calcium transient from the baseline F0 with the following equation: ΔF = (F(t) – F0) / F0. The baseline fluorescence, (F0), was defined as the mean fluorescence (F) from the 2s before the start of the electric shock sequence. We then calculated the peak fluorescence and the area under the curve (integral F/F0) during the whole sequence of electric shocks presentation.

## Quantification and statistical analysis

All statistical analyses were performed using PRISM 10.6.1 (GraphPad Software). All behavioral and imaging data were first tested for normality using the D’Agostino and Pearson normality test. Normally distributed data were analyzed with parametric one-way ANOVA followed by Tukey’s honest significant difference (HSD) post hoc test. For non-Gaussian distributed data, Kruskal-Wallis test was performed followed by Dunn’s multiple comparisons test. Simple linear regression was used to fit a model to some data. We also compared linear regression slopes with Prism (equivalent method to an Analysis of covariance). For the correlation we used parametric Pearson or non-parametric Spearman correlation. Unpaired t-or Mann-Whitney tests were used to compare two group. All graphics were generated with Inkscape 1.3.2 (Inkscape). All statistical comparisons are presented in **Table S2**.

## Acknowledgments

We thank Caleb Larnerd, Stephanie Trouche and Pierre-Yves Plaçais for comments on the manuscript. We acknowledge the imaging facility MRI, member of the France-BioImaging national infrastructure supported by the French National Research Agency (ANR-10-INBS-04, “Investments for the future”). We thank the Montpellier Biocampus *Drosophila* and IPAM facilities (University of Montpellier, CNRS, INSERM, Montpellier, France) for the fly food and the 2-photon microscope, respectively. We thank the Guillaume Cazals at Pole Balar Chemistry, CNRS (Montpellier) for help with the GC/MS. We thank Eloïse Néel for the fly schematic, Nicolas Boucharel for the fly stock maintenance, and Laëtitia Peurien, Théa Michel, Marina Gutiérrez Garcia and Miguel Delgado for preliminary experiments. **Funding:** E.P. is funded by the French Research Agency (ANR-21-CE16-0015-01 and ANR-24-CE92-0014-01), the CNRS and the Institute of Functional Genomics, Montpellier, France. We thank the Bloomington stock center for flies.

## Author contributions

E.P. and N.M. conceived the project with help from C.M. N.M., C.G, C.Z., M.V. and A.-K.T. performed behavioral experiments. N.M analyzed all behavioral experiments. N.M. performed imaging experiments with some help from E.P. N.M. analyzed imaging data. Anatomical data were collected by N.M and P.Z. The manuscript was written by E.P. and N.M with help from C.M.

## Competing interests

The authors declare that they have no competing interests.

## Data, Code and materials availability

All data needed to evaluate the conclusions in the paper are present in the paper and/or the Supplementary Materials. All original reagents presented in this study are available from the Lead Contact upon request.

---

**Fig. S1.**
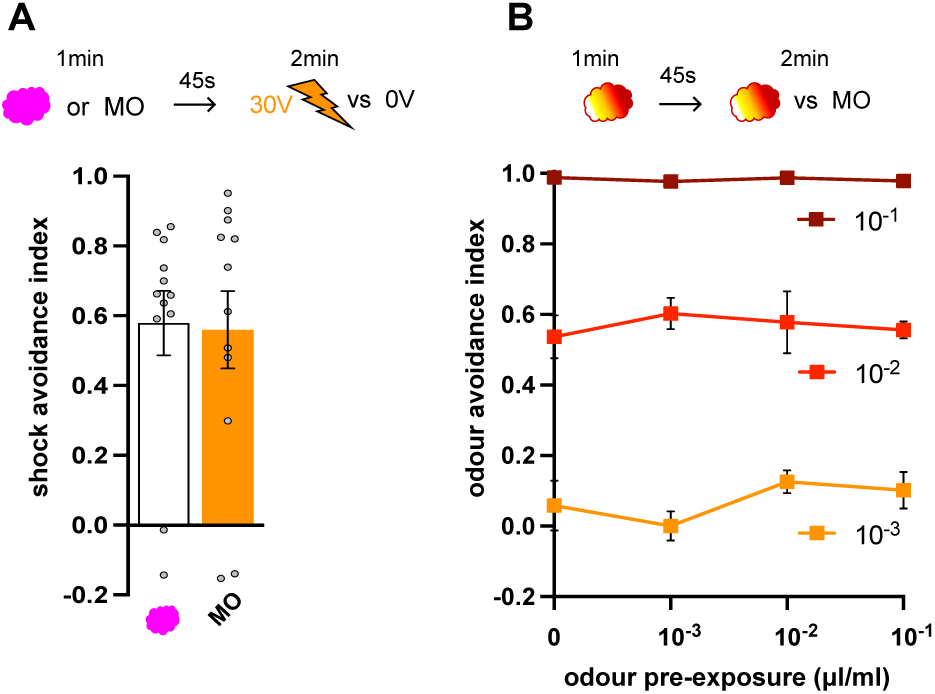
Odor pre-exposure does not alter ensuing shock or odor avoidance. (**A**) Shock avoidance (30V) is not affected by odor pre-exposure (mineral oil (MO) or MCH), n=12. (**B**) Odor avoidance (10^-1^, 10^-2^, 10^-3^ acetophenone vs mineral oil (MO)) is not affected by prior odor pre-exposure (of 0, 10^-1^, 10^-2^, 10^-3^ acetophenone), n=7-11. Top of each graph represents the protocol. Data presented as mean ± standard error of mean (SEM). Individual data points are displayed as dots.

**Fig. S2.**
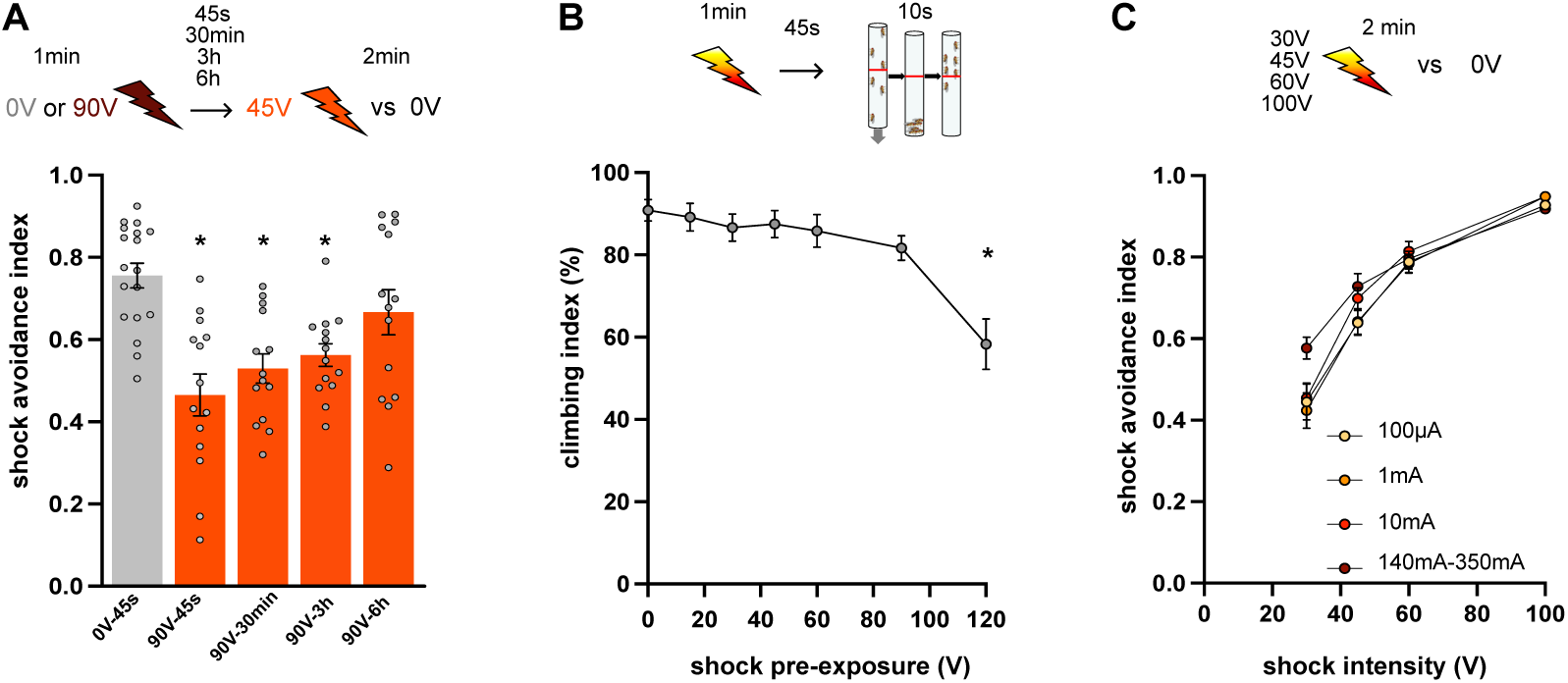
The effect of shock pre-exposure lasts for several hours, does not affect locomotion and is independent of current intensity. (**A**) Shock avoidance (45V) goes back to normal 3 to 6 h after 90V shock pre-exposure, n=14-19. (**B**) Climbing locomotor response is not affected following 0, 15, 30, 45, 60 or 90V shock pre-exposure but is altered following 120 V, n=12. (**C**) Shock avoidance is not affected by any current intensity, n=14. Top of each graph represents the protocol. Data presented as mean ± standard error of mean (SEM). Individual data points are displayed as dots. Asterisks indicate statistically significant differences (p < 0.05).

**Fig. S3.**
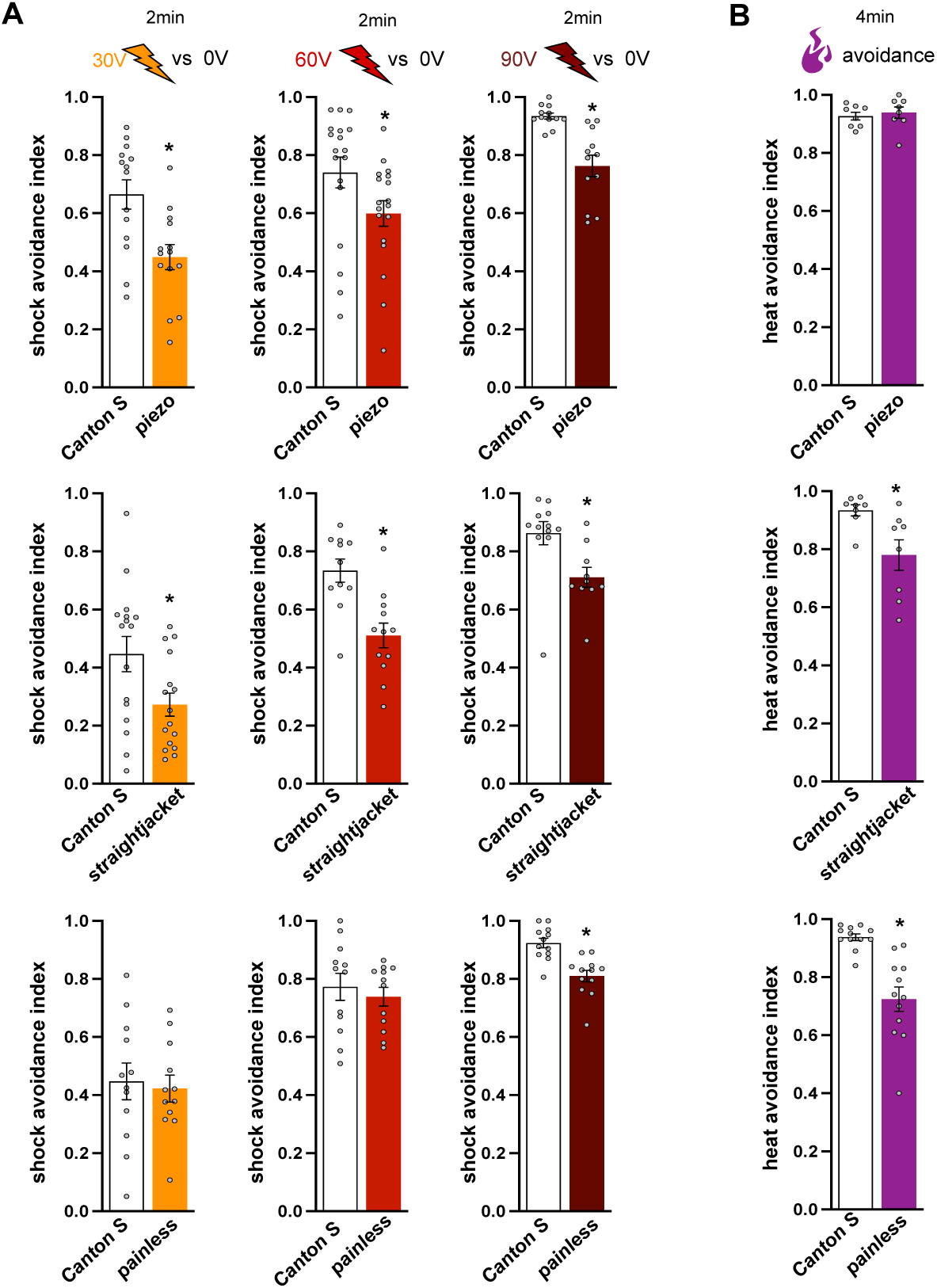
*Piezo, straightjacket* and *painless* mutant are required for shock and heat avoidance. (**A**) 30V, 60V or 90V shock avoidance for *piezo* (top), *straightjacket* (middle) and *painless* (bottom) mutant flies, n=10-18. (**B**) Heat avoidance (42°C) for *piezo* (top), *straightjacket* (middle) and *painless* (bottom) mutant flies, n=8-12. Top of each graph represents the protocol. Data presented as mean ± standard error of mean (SEM). Individual data points are displayed as dots. Asterisks indicate statistically significant differences (p < 0.05).

**Fig. S4.**
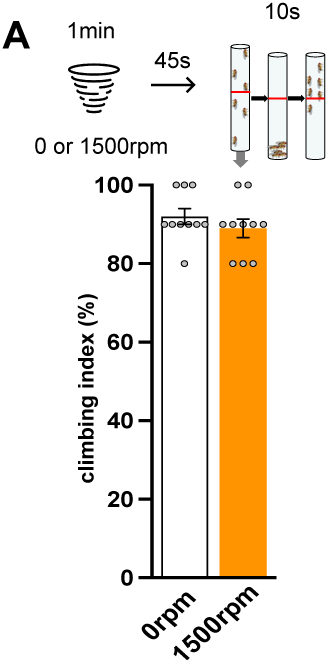
A strong mechanical pre-exposure does not affect climbing behavior. (**A**) Climbing is not affected by prior 1,500 rpm mechanical pre-exposure, n=10. Top of the graph represents the protocol. Data presented as mean ± standard error of mean (SEM). Individual data points are displayed as dots.

**Fig. S5.**
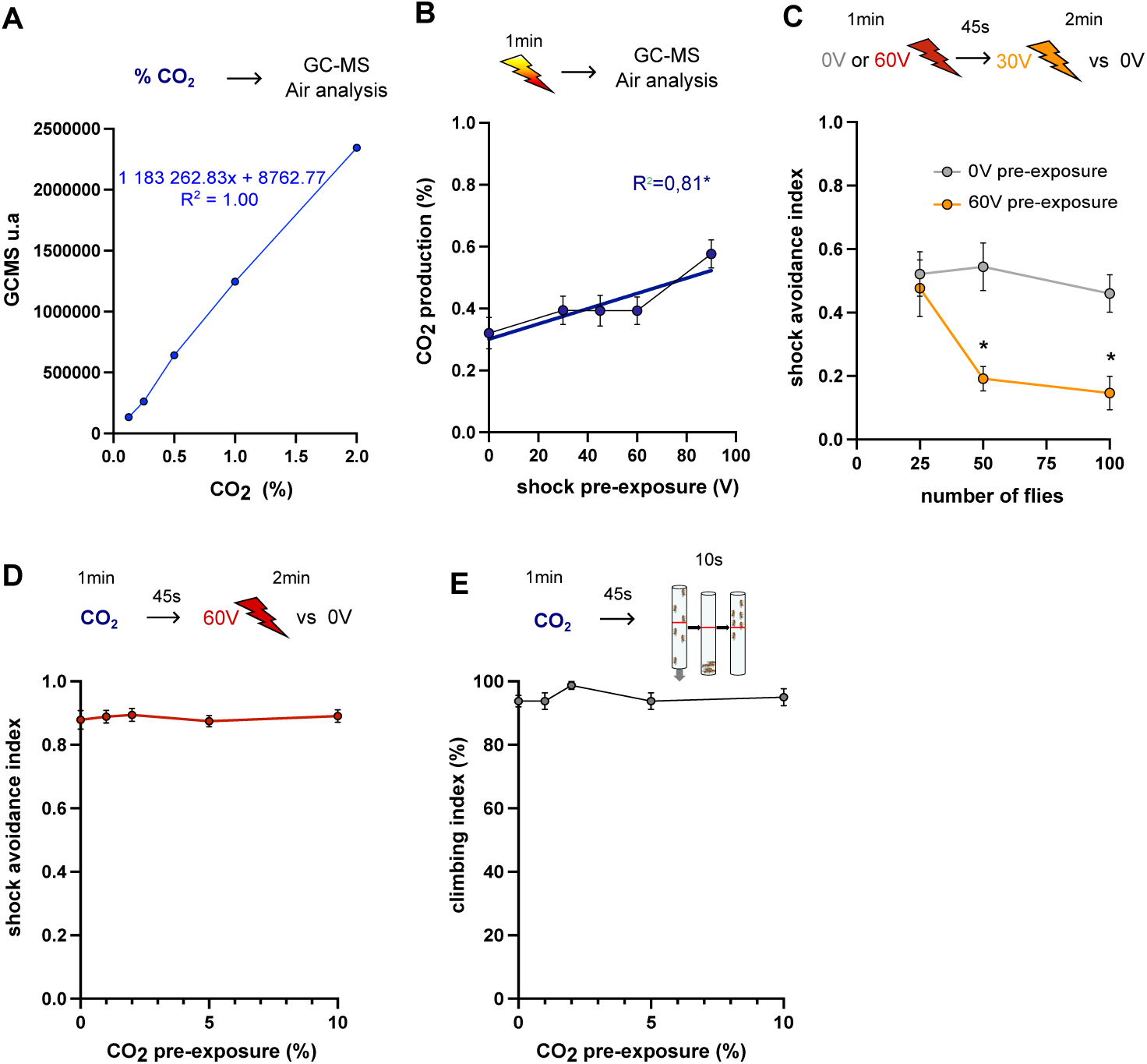
CO_2_ release scales with shock intensity-dependent and does not alter 60V avoidance nor locomotion. (**A**) Mean reference curve between CO_2_ relative abundance measured by GC-MS and real CO_2_ concentration (0,04%, 0,1%, 0,2%, 0,5%, 1% and 2%) injected in GC-MS detector, n=15 (**B**) Fly CO_2_ production depends on the intensity of shock pre-exposure, n=9-11 (**C**) 30V shock avoidance after 60V shock pre-exposure depends on group size (25, 50, 100 flies), n=12. (**D**) 60V shock avoidance is not altered by 1%, 2%, 5%, 10% CO_2_ pre-exposure, n=12. (**E**) Climbing is not affected by 0%, 1%, 2%, 5% or 10% CO_2_ pre-exposure, n=8. Top of each graph represents the protocol. Data presented as mean ± standard error of mean (SEM). Asterisks indicate statistically significant differences (p < 0.05).

**Fig. S6.**
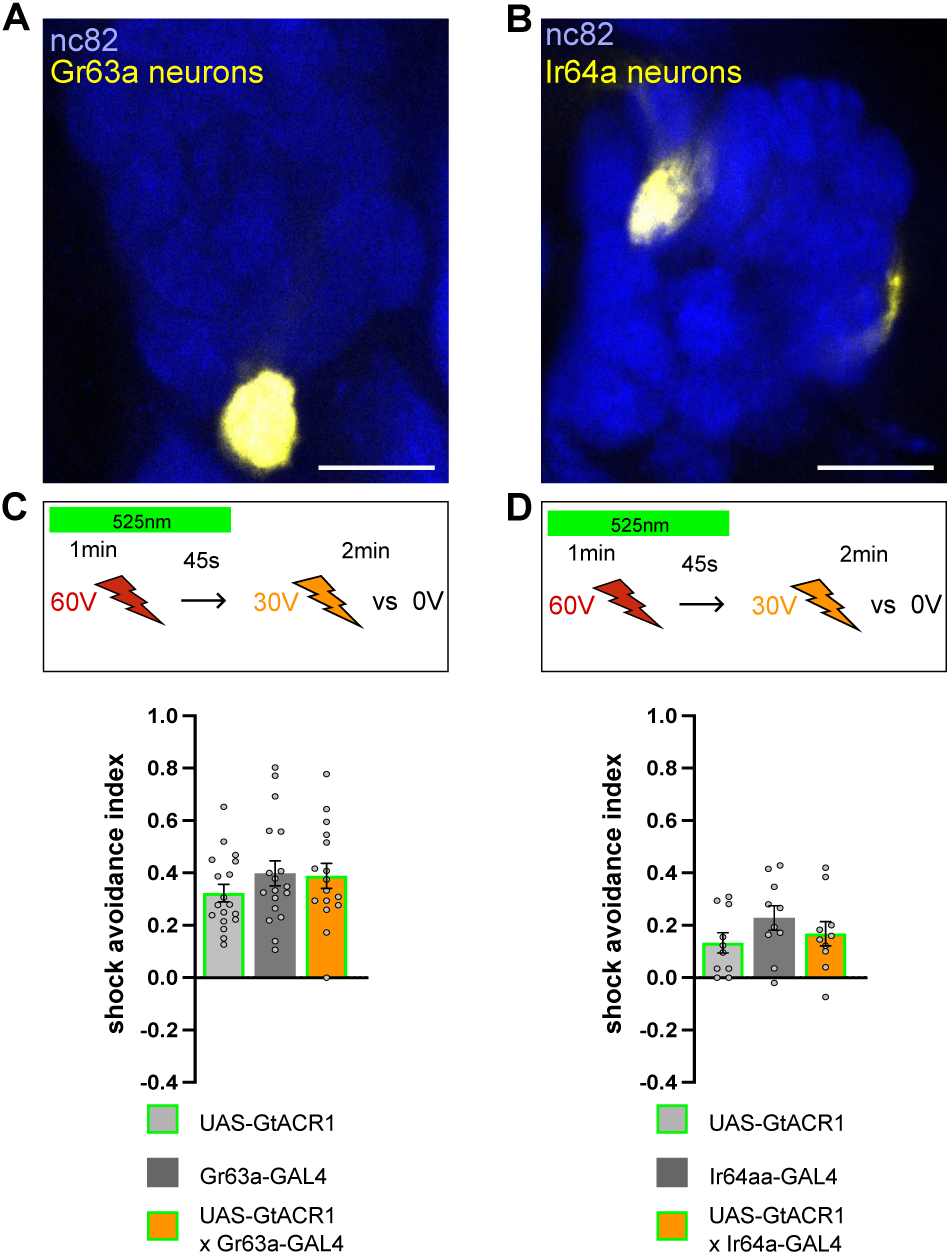
Gr63a-GAL4 and IR64a-GAL4-expressing neurons are not required during 60V shock pre-exposure to modulate subsequent 30V shock avoidance. (**A** and **B**) Gr63a-GAL4 (**A**) and Ir64a-GAL4 (**B**) driving UAS-mCitrine (yellow) targeting CO_2_ sensing neurons innervating the antennal lobe V glomerulus and acid sensing neurons innervating the antennal lobe DC4 glomerulus, respectively. The whole brain is labelled with the presynaptic marker anti-Bruchpilot, nc82 (blue). Scale bar is 20 μm. (**C**) Optogenetic silencing of Gr63a-GAL4 expressing neurons with GtACR1 during 60V shock pre-exposure has no effect on subsequent 30V shock avoidance, n=16-18. (**D**) Optogenetic silencing of Ir64a-GAL4 expressing neurons with GtACR1 during 60V shock pre-exposure has no effect on subsequent 30V shock avoidance, n=8-10. Top of each graph represents the protocol. Data presented as mean ± standard error of mean (SEM). Individual data points are displayed as dots.

**Fig. S7.**
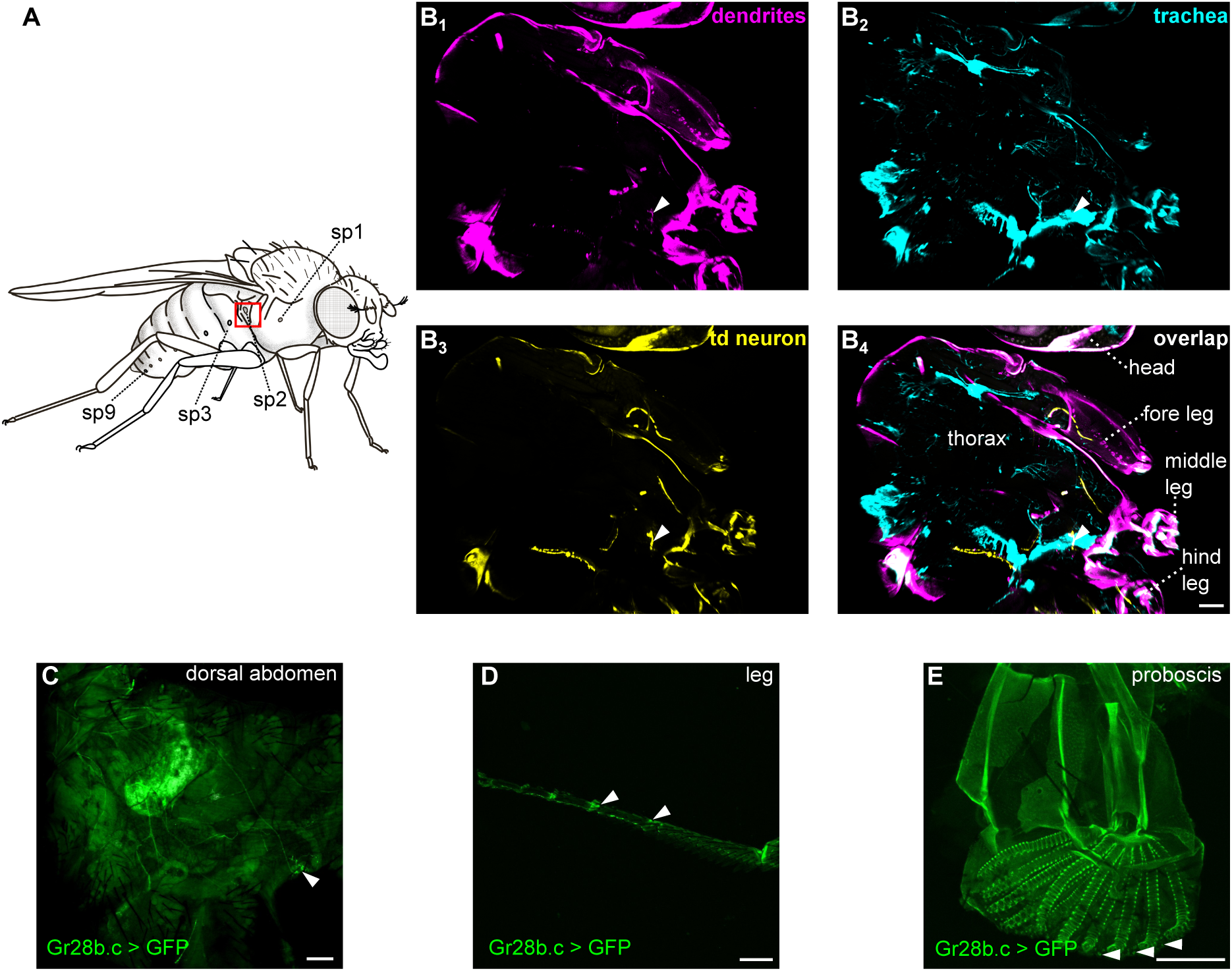
Gr28b.c-GAL4 expression. (**A**) Fly schematic showing the different spiracles on the thorax and abdomen. (**B**) Projection of Gr28b.c-GAL4 expressing neurons in a sagittal section driving UAS-mCitrine (yellow), UAS-DenMark::RFP (magenta) with UV-responding trachea (cyan). The overlap (**B_4_**) shows tracheal (cyan) dendrite (magenta) neurons (yellow) (white arrow). Scale bar is 70 μm. (**C-E**) Gr28b.c-GAL4 driving UAS-mCitrine (green) in the dorsal abdomen (**C**), leg (**D**) and proboscis (**E**) with white arrows showing cell bodies. Scale bar is 100 μm.

**Fig. S8.**
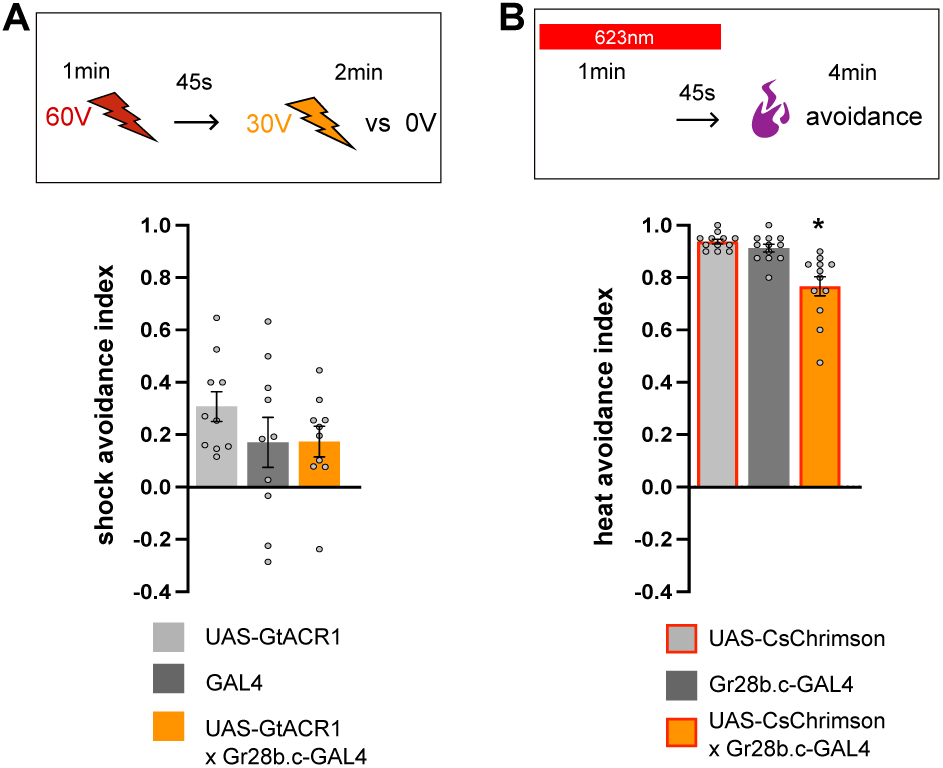
Gr28b.c-expressing (td) neurons activation affects heat avoidance. (**A**) Control experiment for Gr28b.c-expressing (td) neurons with GtACR1 without optogenetic stimulation for 60V shock pre-exposure before 30V shock avoidance, n=10. (**B**) Optogenetic activation of Gr28b.c expressing (td) neurons with csChrimson during a 1 min pre-exposure significantly reduces subsequent heat avoidance, n=10-12. Top of each graph represents the protocol. Data presented as mean ± standard error of mean (SEM). Individual data points are displayed as dots. Asterisk indicates statistically significant differences (p < 0.05).

**Fig. S9.**
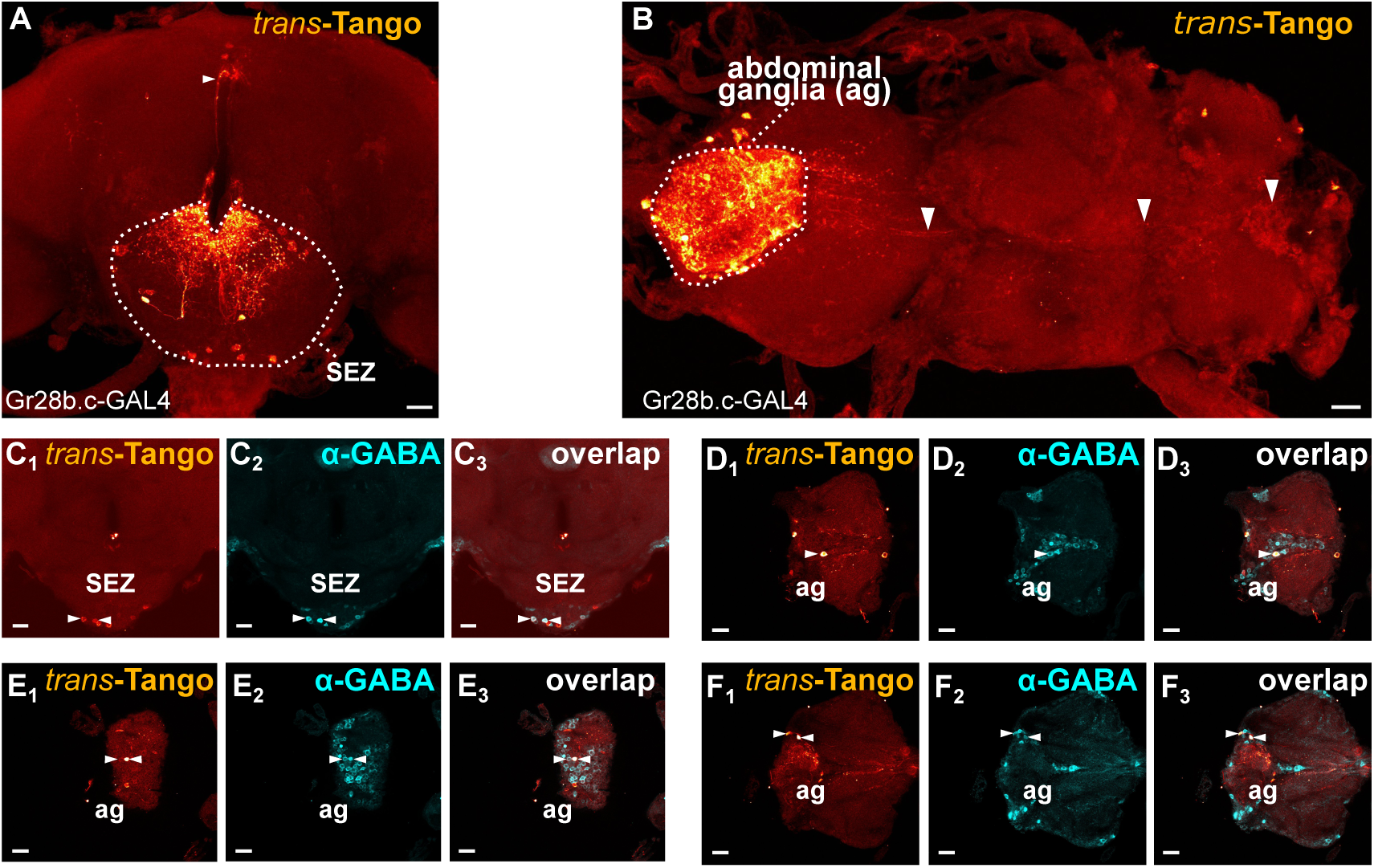
Downstream targets of Gr28b.c-expressing neurons. (**A**-**B**) Expression of the *trans*-Tango signal of Gr28b.c-GAL4 reveals post-synaptic neurons (red hot) of td neurons expressed in the brain (**A**), mostly in the SEZ, with some neurons targeting the SMP region (white arrow), and expressed in the VNC (**B**), mostly in the abdominal ganglia with ascending neurons (white arrows). Scale bar is 20 μm. (**C**-**F**) GABAergic staining of the Gr28b.c *trans*-Tango showing GABA+ signal in the ZEZ (**C**) and VNC (**D**, **E** and **F**) cell bodies. Scale bar is 20 μm.

**Fig. S10.**
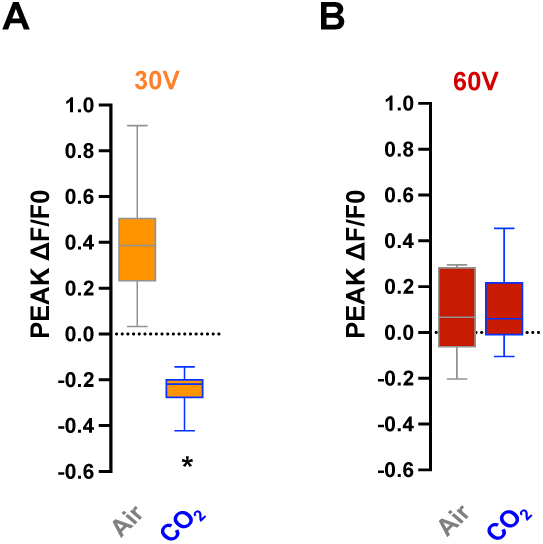
Pre-exposing flies to CO_2_ leads to decreased 30V but not for 60V dopaminergic neurons shock response. (**A**) Peak delta F/F0 calcium transients ± standard error of the mean (SEM) measured from PPL1-γ1pedc targeted by MB504B-GAL4 to 30 V shocks significantly decreases after 10% CO_2_ pre-exposure, n=11. (**B**) Peak delta F/F0 calcium transients ± standard error of the mean (SEM) measured from PPL1-γ1pedc targeted by MB504B-GAL4 to 60 V shocks is not affected by 10% CO_2_ pre-exposure, n=7-8. Data presented as boxplot with the min, median and max value. Asterisk indicates statistically significant differences (p < 0.05).

**Fig. S11.**
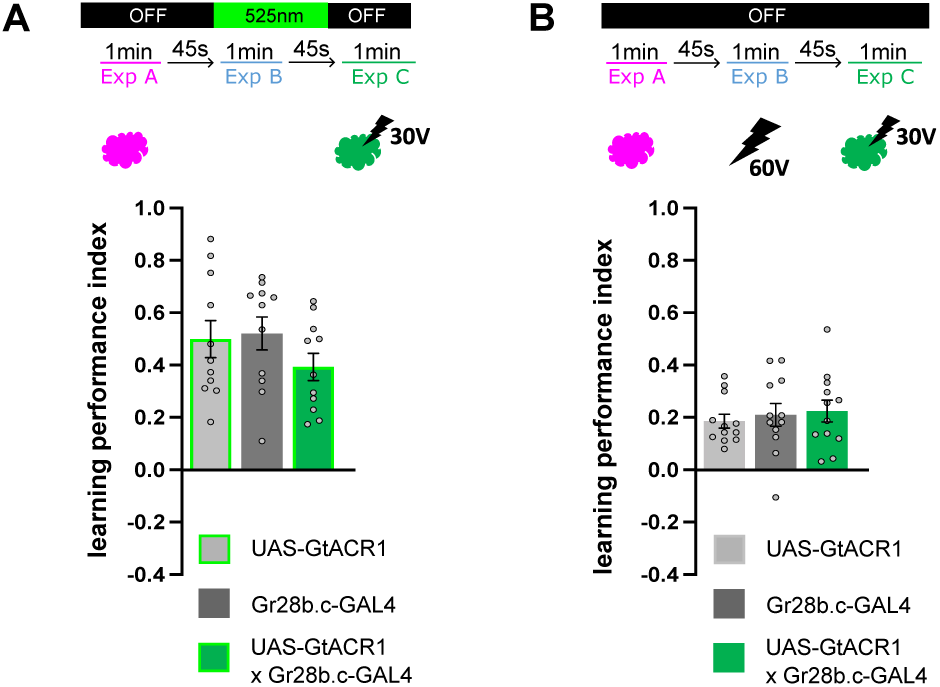
Optogenetic inhibition of Gr28b.c-expressing (td) does not alter subsequent odor C+30V shock pairing. (**A**) Optogenetic inhibition of Gr28b.c-expressing (td) neurons driving GtACR1 during 0V pre-exposure does not affect memory retrieval of odor A vs C, n=12. (**B**) Control experiment for Gr28b.c-expressing (td) neurons with GtACR1 without optogenetic stimulation during 60V pre-exposure before learning odor C + 30V, n=12. Top of each graph represents the protocol. Data presented as mean ± standard error of mean (SEM). Individual data points are displayed as dots.

**Table S1.** All resources information needed to perform all experiments and analyses.

**Table S2.** All statistical comparisons for all figure panels.

## Legend of Movies

**Movie S1.** Downstream targets of Gr28b.c-expressing neurons. *trans*-Tango signal in the brain

**Movie S2.** Downstream targets of Gr28b.c-expressing neurons. *trans*-Tango signal in the VNC.

## Notes

### Competing Interest Statement

The authors have declared no competing interest.

